# Apple scar skin viroid disease induced physicochemical and metabolic alterations in Apples

**DOI:** 10.64898/2026.07.28.741181

**Authors:** Prem Chand, Hema Kumari, Esha Devi, Rishav Kumar, Santosh Watpade, Shyam Kumar Masakapalli

## Abstract

Apple scar skin disease (ASSD), caused by Apple scar skin viroid (ASSVd), is characterized by peel scarring, cracking, dappling, and fruit deformation, resulting in reduced fruit quality and marketability. Despite its economic importance, the physicochemical and metabolic alterations underlying disease progression remain poorly understood. To address this knowledge gap, apple fruits representing four stages of ASSD (healthy, lightly infected, moderately infected, and highly infected) were comprehensively characterized. ASSVd infection was confirmed by RT-PCR, amplicon sequencing, and phylogenetic analysis. Fruit morphology and quality attributes, including firmness, total soluble solids (TSS), pH, titratable acidity (TA), and total phenolic content (TPC), were evaluated. ASSVd infection significantly reduced fruit weight and firmness and altered TSS and TA, indicating progressive deterioration of fruit quality. To investigate the underlying metabolic changes, peel and pulp tissues were analysed separately using gas chromatography-mass spectrometry (GC-MS), while major soluble sugars were quantified by ^1^H nuclear magnetic resonance (^1^H NMR) spectroscopy. Integrated metabolomic analyses revealed distinct tissue-specific metabolic reprogramming during disease progression. Major soluble sugars declined significantly during early infection, followed by tissue-dependent recovery at later stages, whereas organic acids, amino acids, phenolics, lipids, polyols, and pentacyclic triterpenoids exhibited dynamic stage-dependent changes. Notably, lupeol accumulated progressively, whereas ursolic acid and oleanolic acid declined, indicating disease-associated alterations in host triterpenoid metabolism. Multivariate analyses demonstrated clear metabolic separation among disease stages. Lupeol, ursolic acid, and chlorogenic acid were identified as candidate discriminatory metabolites in the peel, whereas myo-inositol, chlorogenic acid, and aspartic acid were identified in the pulp. Collectively, these findings demonstrate that ASSD induces coordinated, tissue-specific physicochemical and metabolic reprogramming that compromises postharvest fruit quality and reshapes defence-associated metabolism. This study provides the first integrated metabolomic characterization of ASSD progression and identifies potential metabolic biomarkers for disease diagnosis and severity assessment.

**Graphical Abstract:** 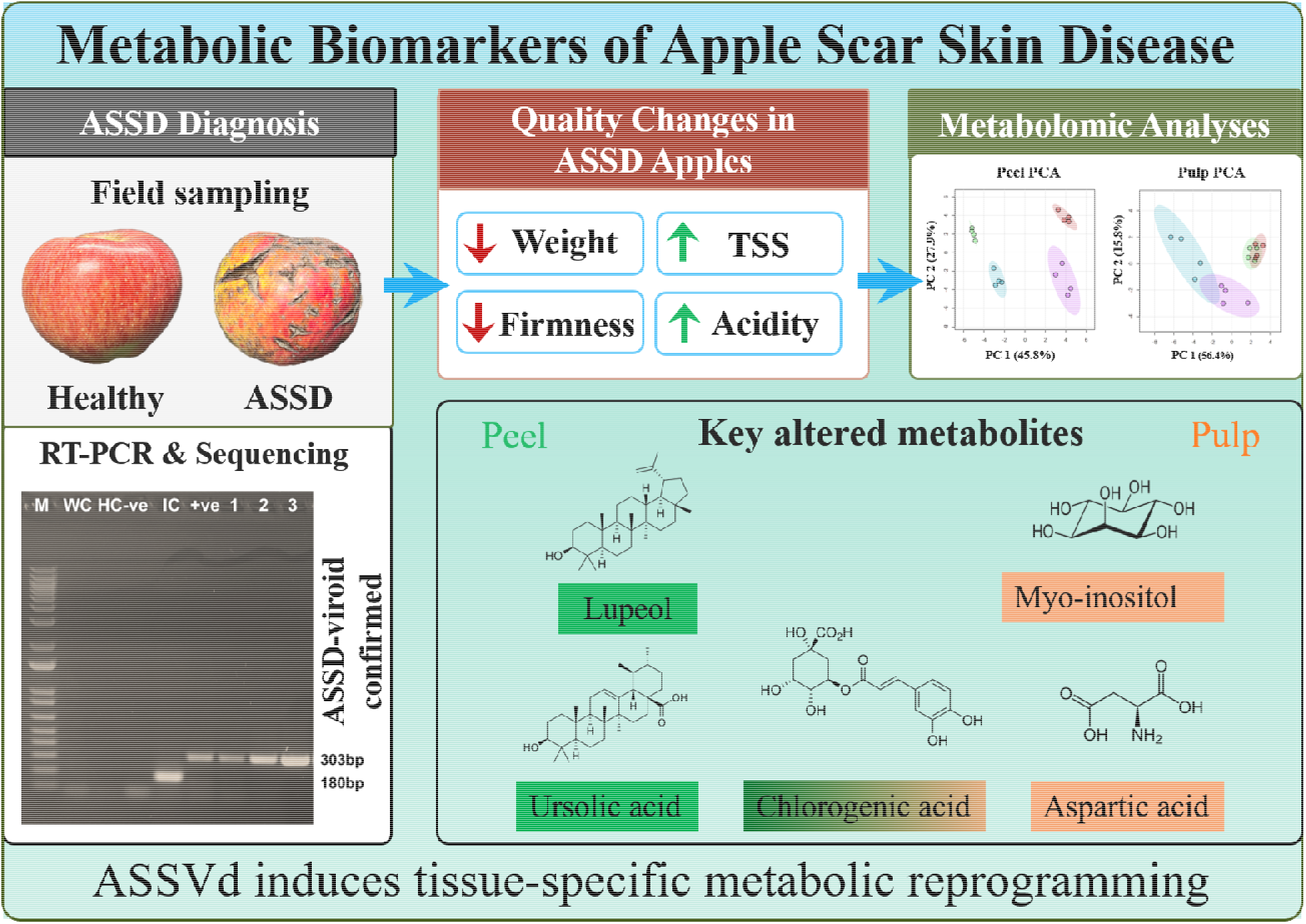

## 1. Introduction

Apple (*Malus × domestica* Borkh.) is one of the most economically important temperate fruit crops and is known for its nutritional and phytochemical composition (Boyer *et al*., 2004; Wani *et al*., 2018; Dadwal *et al*., 2023). Global apple production is dominated by China (47.0 Mt), European Union (11.52 Mt), United States (5.0 Mt), Türkiye (1.9 Mt), and India (2.6 Mt) (FAOSTAT, 2025a). In India, apple is cultivated on approximately 0.304 million hectares, concentrated mainly across the Himalayan region, particularly in Jammu and Kashmir, Himachal Pradesh, and Uttarakhand, where it provides an important source of income for small and marginal farmers (Kumar *et al*., 2016; Agricoop, 2025). Despite accounting for approximately 6.58% of the global area under apple cultivation, India contributes only about 3.2% of global production (FAOSTAT, 2025b). Low productivity has been associated with several factors, including traditional orchard management, limited varietal diversification, inadequate winter chilling, and susceptibility to biotic and abiotic stresses (Pramanick *et al*., 2015).

Among the biotic constraints affecting apple production, diseases caused by viruses and viroids are particularly difficult to manage because infected perennial trees can serve as long-term sources of inoculum. Apple scar skin disease (ASSD), associated with Apple scar skin viroid (ASSVd), primarily affects fruit appearance and can substantially reduce marketability (Sharma *et al*., 2020). ASSVd belongs to the genus *Apscaviroid* within the family *Pospiviroidae* and was among the first viroids that has been documented to infect pome fruit species, including *Malus*, *Pyrus*, and *Cydonia* spp. (Hashimoto and Koganezawa, 1982). Unlike many pathogens that produce conspicuous symptoms on vegetative tissues, ASSVd infection may remain visually undetected in leaves and shoots while causing severe scarring, cracking, and discoloration of the fruit surface (Hadidi *et al*., 2024; Sharma *et al*., 2020; Vadamalai *et al*., 2024). ASSD is of particular concern in apple-growing regions of India. In some orchards of Himachal Pradesh, ASSVd incidence has been reported in up to 90% of trees, with an estimated 72.68% reduction in net return per hectare (Sharma *et al*., 2020).

Although the external symptoms and economic consequences of the ASSD have been documented (Desvignes *et al*., 1999; He *et al*., 2026; Tian *et al*., 2022; Sharma *et al*., 2020), much less is known about the biochemical perturbations occurring in fruit composition as symptoms progress. Studies including metabolic consequences of ASSVd infection in apples has emerged only recently. In young apple saplings, Li *et al*. (2023) reported that ASSVd impaired vegetative growth by reducing leaf size, photosynthetic performance, and mineral nutrient accumulation, accompanied by alterations in metabolites related to phenylpropanoid, isoflavonoid, and carbohydrate metabolism. In fruit, He *et al*. (2026) showed that ASSVd affected phenylpropanoid pathway, suppressing anthocyanin biosynthesis in dapple fruit, while promoting suberin biosynthesis in scar skin fruit. Using an apple callus infection system, He *et al*. (2026) further demonstrated that ASSVd downregulated key genes involved in anthocyanin biosynthesis, resulting in reduced pigment accumulation. Despite these advances, current knowledge is largely confined to specific tissues or individual metabolic pathways, and a comprehensive understanding of the physicochemical and tissue-specific metabolic alterations that occur during successive stages of ASSD symptom development in apple fruit is still lacking. Metabolite profiling provides a powerful approach to address this knowledge gap by linking visible disease progression with changes in fruit biochemistry. Gas chromatography-mass spectrometry (GC-MS) provides broad coverage of primary and secondary metabolites (Salem *et al*., 2020; Lisec *et al*., 2006), whereas ^1^H nuclear magnetic resonance (^1^H NMR) spectroscopy enables robust and reproducible detection and quantification of abundant metabolites in biological samples (Tian *et al*., 2016). When integrated with morphological and physicochemical assessments, GC-MS and ^1^H NMR metabolomics enable comprehensive characterization of tissue-specific metabolic responses to ASSVd across successive stages of symptom development. Therefore, this study aimed to comprehensively characterize the physicochemical and tissue-specific metabolic alterations associated with ASSD progression in apple fruit and to identify key metabolites associated with successive stages of ASSVd infection.

## 2. Materials and methods

All the chemicals and reagents were purchased from Sigma-Aldrich, unless mentioned in the methods.

### 2.1 Plant Material

Apple fruits showing characteristic symptoms of apple scar skin disease (ASSD), together with asymptomatic fruits, were collected from farmers’ orchards in Chithala village, Kotkhai tehsil, Shimla district, Himachal Pradesh, India (31°06′11.8″ N, 77°32′10.0″ E), in September 2024. Based on the extent of visible symptoms including percentage of fruit surface affected, type of scarring, cracking, and pigmentation, the apple fruits were classified into four groups: healthy (H), lightly infected (LI), moderately infected (MI), and highly infected (HI). The fruits were thoroughly washed with distilled water, surface-dried with muslin cloth, and stored at 4 °C until further analysis. For metabolite analysis, peel and pulp tissues were separated and immediately frozen in liquid nitrogen. The tissues were subsequently ground to a fine powder, freeze-dried and stored at −80 °C until further use.

### 2.2 Molecular characterization

Total RNA was isolated from fruit samples using the Spectrum™ Plant Total RNA Kit (Sigm-Aldrich, USA) following the manufacturer’s instructions. RNA was quantified using a Thermo Scientific NanoDrop 2000. cDNA was synthesized from 1 µg RNA using the RevertAid™ cDNA Synthesis Kit (Thermo Fisher Scientific Inc.) as per the manufacturer’s protocol. All reactions were set up under ice-cold conditions to prevent premature cDNA synthesis and RNA degradation. The reaction was incubated at 25 °C for 5 min, 42 °C for 60 min, and 75 °C for 10 min. The synthesized cDNA was stored at −20 °C and used for PCR amplification. The RT-PCR reaction mixture comprised 12.5 μl of EmeraldAmp GT PCR Master Mix (Takara Bio), 10 μM of each primer, 2.0 μl of cDNA, and nuclease-free water to a final volume of 25 μl. Primers designed by Sharma *et al*. (2020) and RT-PCR conditions optimized by Watpade *et al*. (2021) were used. Gel extraction was carried out using the QIAquick® Gel Extraction Kit (Qiagen), and the purified amplicon was subsequently sequenced (HiMedia Laboratories Pvt. Ltd., India).

### 2.3 Morphometric Measurements

Fruit weight was measured using an analytical balance (ME204, Mettler Toledo, Switzerland; 0.1 mg readability) and expressed in grams. Fruit height (polar diameter) and width (equatorial diameter) were measured with a digital Vernier caliper (CD-6″ CSX, Mitutoyo, Japan). The fruit shape index was expressed as the ratio of fruit height to equatorial diameter.

### 2.4 Firmness Measurement

Fruit firmness was measured at two opposite positions along the equatorial region after removing a small section of peel. Measurements were made using a digital fruit pressure tester (ACSY4, Acucal, India) fitted with an 11 mm cylindrical stainless-steel probe. Firmness was expressed as kilogram-force (kgf). For each fruit, the mean of the two measurements was calculated, and ten fruits per infection stage were used for statistical analysis.

### 2.5 Total Soluble Solids (TSS)

Total soluble solids (TSS) were measured in freshly extracted apple juice using a handheld refractometer (ACSRF-1, Acucal, India). The instrument was calibrated with distilled water before measurement, and the results were expressed as °Brix at 20 °C.

### 2.6 pH Measurement

Fresh apple pulp (5 g) was homogenized with 10 mL of distilled water using a mortar and pestle and centrifuged at 14,000 rpm at 20 °C for 5 minutes (Dadwal *et al*., 2023). The pH of the supernatant was measured using a SevenCompact™ S220 pH meter (Mettler Toledo, Switzerland).

### 2.7 Titratable Acidity (TA)

Titratable acidity (TA) was determined according to Dadwal *et al*. (2023). Apple juice (10 mL) was titrated against 0.1 N NaOH until a persistent pale pink endpoint was reached using phenolphthalein as the indicator. TA was expressed as gram malic acid equivalents L^-1^(g/L MAE) of juice.

### 2.8 Determination of Total Phenolic Content

Total phenolic content (TPC) was determined using the Folin-Ciocalteu assay following Kupina *et al*. (2019), with minor modifications. Lyophilized peel or pulp (10 mg) was extracted with 1 mL of 70% (v/v) acetone and centrifuged at 10,000 rpm for 15 min at 4 °C. An aliquot (100 µL) of the supernatant was mixed with 500 µL of 10% (v/v) Folin-Ciocalteu reagent and incubated for 5 min. Subsequently, 400 µL of 5% (w/v) sodium carbonate solution was added, and the reaction mixture was incubated in the dark at room temperature for 20 min. Absorbance was measured at 765 nm using a spectrophotometer. Total phenolic content was quantified from a gallic acid standard curve and expressed as mg gallic acid equivalents (GAE) g^-1^ dry weight (DW).

### 2.9 Metabolite extraction for ^1^HNMR and GC-MS analysis

Lyophilized peel and pulp samples (50 mg DW) were extracted with 1 mL of water:chloroform:methanol (1:1:3, v/v/v), based on Lisec *et al*. (2006). For GC-MS analysis, 50 µL of ribitol (0.01%, w/v) was added as an internal standard. Samples were sonicated for 5 min in ultrasonic bath, followed by heating at 65-70 °C for 3-5 min and cooled on ice for 2-3 min. The heating and cooling extraction cycle was repeated three times to enhance metabolite recovery. Finally, the samples were incubated in thermoshaker at 70 °C, 900rpm for 5 min. The extracts were vortexed and centrifuged at 13,000 rpm for 10 min. For ^1^H-NMR analysis, 200 µL of the supernatant was vacuum dried and redissolved in 600 µL of D_2_O containing 0.1 mM phosphate buffer (pH 7.2) and 0.01% (w/v) trimethylsilylpropanoic acid (TSP). The solution was transferred to 5 mm NMR tubes for analysis. For GC-MS analysis, 50 µL of the supernatant was vacuum dried. The dried residue was derivatized according to the procedure described by Masakapalli *et al*., (2014). The sample was methoxylated by adding 35 µL of methoxyamine hydrochloride (20 mg mL^-1^) in pyridine, followed by incubation at 37 °C for 2 hours. Subsequently, 49 µL of N-methyl-N-(trimethylsilyl)trifluoroacetamide (MSTFA) was added and samples were incubated for 30 min for silylation. The derivatized samples were centrifuged at 14000 rpm for 10 min and transferred into glass inserts for GC-MS analysis.

### 2.10 ^1^H-NMR data acquisition and quantification

^1^H NMR spectra were acquired using a 500 MHz NMR spectrometer (JEOL ECX) at 298 K using a one-dimensional proton experiment with water suppression. A 90° pulse angle and a relaxation delay of 10 s were applied to ensure quantitative reliability. Each spectrum consisted of 64 scans. Spectra were processed using JEOL Delta (Version 6.1.0) software. Phase and baseline corrections were applied uniformly, and TSP (δ 0.00 ppm) was used as the chemical shift reference. Sugars were identified using analytical standards and reported literature chemical shifts. Calibration curves for the quantification of soluble sugars were established using standard solutions with concentrations ranging from 0.8 to 13.3 mg mL^-1^. Quantification was based on well-resolved proton resonances of α-glucose (δ 5.22-5.24 ppm), β-glucose (δ 4.64-4.66 ppm), fructose (δ 4.09-4.11 ppm), and sucrose (δ 5.40-5.42 ppm). Concentrations were determined using external calibration curves (R² ≥ 0.99) based on TSP-normalized integrals. Final sugar concentrations were expressed as mg g^-1^ dry weight.

### 2.11 GC-MS data acquisition and processing

GC-MS analysis was performed using a GC-MS 5977B system (Agilent Technologies) equipped with an HP-5ms capillary column (30 m × 250 μm × 0.25 μm). The system was operated in splitless mode, with helium as the carrier gas at a constant flow rate of 0.6 mL min^-1^. The oven temperature was initially maintained at 50 °C for 1 min, increased to 200 °C at 10 °C min^-1^ and held for 4 min, and then raised to 300 °C at 5 °C min^-1^, with a final hold of 10 min. Raw data were processed using MetAlign software for baseline correction and peak alignment. Metabolite identification was performed using Agilent MassHunter Qualitative Navigator software, based on retention time and mass fragmentation patterns with reference to the NIST library (v2.3, 2017). Only metabolites with identification scores ≥70% were retained. Peak abundances were normalized to the internal standard-Ribitol and used for downstream statistical analyses including fold-change calculations and multivariate analysis.

### 2.12 Statistical and multivariate data analysis

Data are expressed as the mean ± standard deviation (SD), unless otherwise specified. Morphometric and physicochemical measurements were based on ten biological replicates per infection stage, whereas biochemical and metabolomic analyses were performed using four biological replicates per infection stage. For the morphometric and physicochemical traits presented in Figure. 1 (fruit height, weight, firmness, total soluble solids, pH, and titratable acidity), total phenolic content presented in Figure. 2, and individual sugars presented in Figure. 3, differences among infection stages were assessed by one-way ANOVA, followed by Dunnett’s post hoc test, with each infected group compared against the healthy control. For the combined comparison of total soluble solids and individual sugars presented in Figure. 4, all pairwise comparisons were performed using one-way ANOVA followed by Tukey’s HSD post hoc test among infection stages. A *p*-value < 0.05 was considered statistically significant. Unless otherwise stated, statistical significance in all figures was denoted as follows: *p* < 0.05 (*), *p* < 0.01 (**), *p* < 0.001 (***), *p* < 0.0001 (****); ns, not significant. For GC-MS data, peak abundances were first normalized to the ribitol (internal standard). The resulting dataset was median-normalized and log□-transformed, followed by Pareto scaling for multivariate analysis. Metabolites differing among infection stages were identified separately for peel and pulp using one-way ANOVA followed by FDR correction for multiple comparisons. Metabolites with an FDR-adjusted *p*-value < 0.05 were considered statistically significant. For visualization of perturbations relative to healthy fruits, log□ fold change (log□FC) was determined as the difference in mean values between log□-transformed abundance of each infected group and that of the corresponding healthy group. Candidate discriminatory metabolites were prioritized by integrating ROC performance, PLS-DA VIP scores, and statistical significance across infection stages. Statistical and metabolomic analyses were performed using MetaboAnalyst 6.0, GraphPad Prism 8, RStudio, and Microsoft Excel. GC-MS and NMR data were processed using MSD ChemStation (vF.01.03.2357), Agilent MassHunter (vB.08.00), and JEOL Delta (v6.1), respectively.

**Figure 1.**
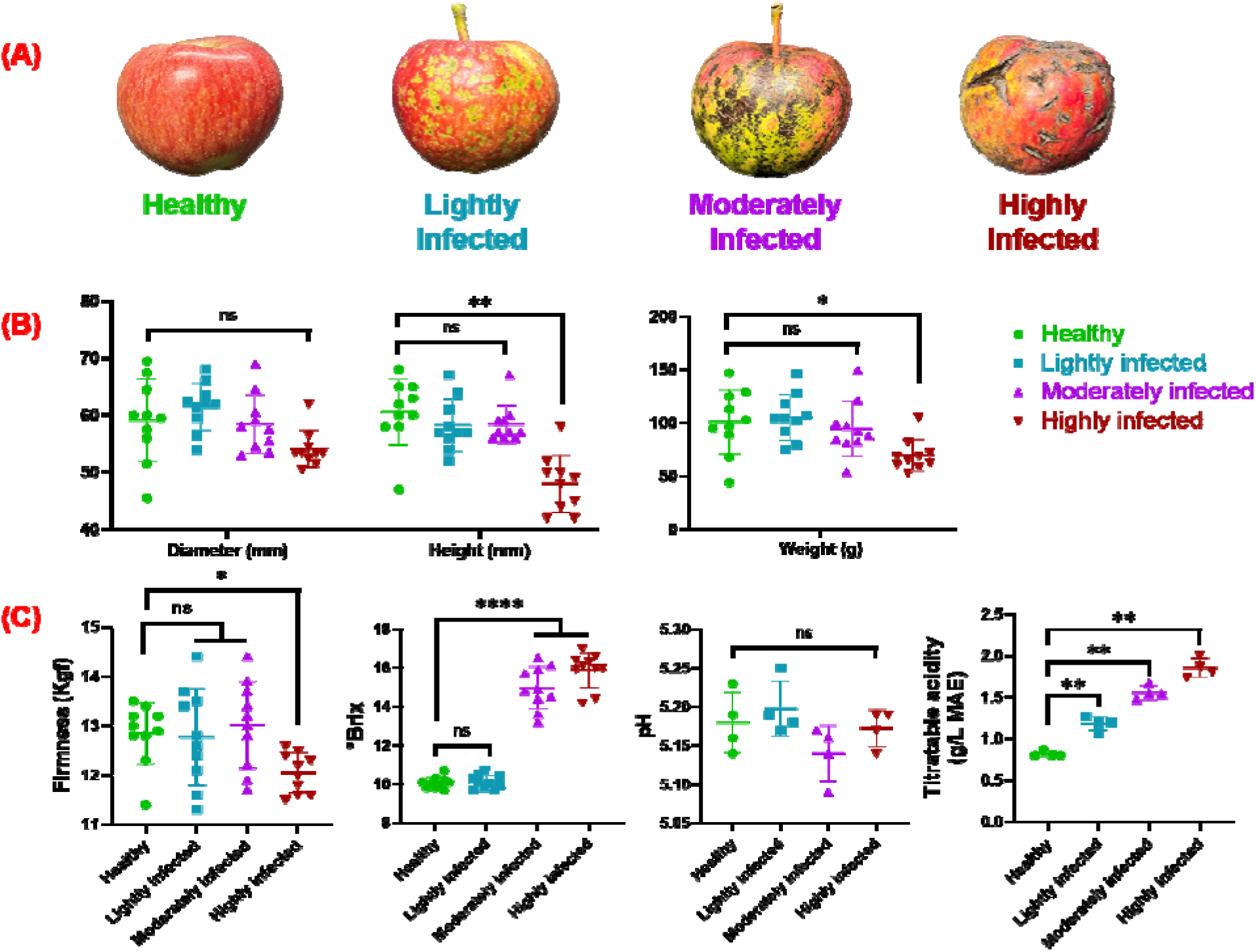
Morphological and physicochemical characteristics of apple fruits at different stages of apple scar skin disease (ASSD). **(A)** Representative fruits classified as healthy, lightly infected, moderately infected, and highly infected according to visible symptom severity. **(B)** Fruit diameter, height, and weight and **(C)** firmness, total soluble solids (°Brix), pH, and titratable acidity (g/L malic acid equivalent) across the disease stages. (*n* = 10 biological replicates per stage).

## 3. Results

### 3.1 Disease stratification and molecular validation

Apple fruits were classified as healthy, lightly infected, moderately infected, or highly infected according to the severity of visible symptoms. Healthy fruits had a smooth peel and uniform appearance, whereas infected fruits showed increasing surface discoloration, scarring, and cracking with symptom severity (Figure. 1A). RT-PCR produced an amplicon of approximately 303 bp in samples from all three symptomatic groups, whereas no amplification was detected in healthy fruits (Supplementary Figure. S1). Sequence analysis showed that the ASSVd sequences obtained from the three symptomatic groups matched previously reported ASSVd isolates. The sequence from the highly infected (HI) group showed 100% identity with the ASSVd-C isolate Liaoning (AY972082.1), whereas the sequences from the moderately infected (MI) and lightly infected (LI) groups showed 100% identity with *Apscaviroid cicatricimali* isolate MS4-5 (KY963664.1) and Apple scar skin viroid isolate Bong Hwa (KP765428.1), respectively. Phylogenetic analysis placed all three sequences within the broader ASSVd lineage, although they occupied different positions within the tree (Supplementary Figure. S2).

### 3.2 Morphological and quality changes in apple fruit during ASSD progression

Morphological parameters were evaluated to determine the effects of ASSD progression on apple fruit morphology. Fruit diameter showed no significant difference between healthy and infected samples (Figure. 1B). Fruit height and weight were observed to be lower in highly infected fruits, with 20.8% and 30.9%, reduction respectively, compared to control. Quality attributes serve as indicators of fruit physicochemical traits, including firmness, TSS, pH, and TA, exhibited significant changes with increasing ASSD severity. Although, lightly and moderately infected fruits showed no significant differences in the firmness, the highly infected fruit reduced by 6.7%. Similarly, no difference in TSS was observed between healthy and lightly infected fruits, however the values increased by 48.7% and 57.9% in moderately and highly infected fruits, respectively. Fruit pH showed no significant differences among infection stages. In contrast, TA increased with symptom severity, at 44.8%, 89.8%, and 126.5% in lightly, moderately, and highly infected fruits, respectively, compared with healthy fruits (Figure. 1C).

### 3.3 ASSD-induced changes in total phenolic content (TPC) of apple peel and pulp

TPC differed markedly between peel (0.67-1.3 mg GAE g^-1^ DW) and pulp (0.03-0.07 mg GAE g^-1^ DW) throughout disease progression (Figure. 2). Across the four stages, peel contained on an average 19.7-fold more phenolics than pulp. Relative to healthy peel, TPC increased by 1.37 and 1.42-fold in lightly and moderately infected fruits, respectively, whereas 20.3% reduction was observed in highly infected fruits. In pulp, TPC remained low and did not differ significantly among infection stages.

**Figure 2.**
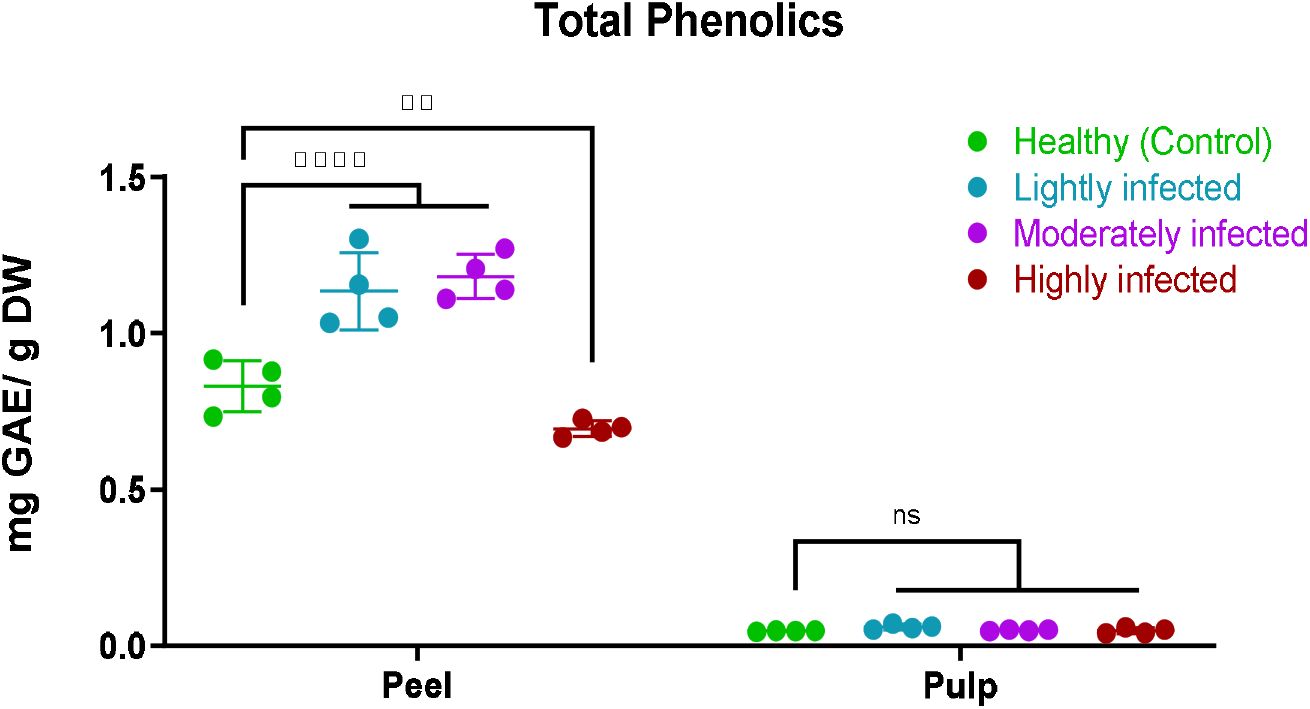
Total phenolic content (TPC) of apple peel and pulp at different stages of apple scar skin disease (ASSD). TPC is expressed as mg GAE g^-1^ DW. (*n* = 4 biological replicates per stage).

### 3.4 Tissue-specific quantitative profiling of soluble sugars by ^1^H NMR

Quantitative ^1^H-NMR analysis showed significant differences in major soluble sugars (fructose, glucose, and sucrose) across infection stages in both peel and pulp (Figure. 3). Fructose concentrations declined markedly during the early stages of ASSD (54.8 mg/g to14.0 mg/g in peel and 141.8 mg/g to 43.8 mg/g in pulp). In peel, fructose decreased by 74.4% in lightly infected fruits compared with healthy fruits (p < 0.0001) and remained 56.47% lower in moderately infected fruits. However, in highly infected peel, fructose was only 3.83% lower than in healthy fruits, with no significant difference. In pulp, fructose decreased by 69.09% in lightly infected fruits (p < 0.0001), followed by reductions of 54.28% and 24.79% in moderately and highly infected fruits, respectively. Although fructose levels partially recovered with increasing disease severity, the reduction remained significant in highly infected pulp (p < 0.0001). Glucose exhibited a trend similar to that of fructose, with a pronounced decline during the early stages of ASSD followed by recovery at later stages (60.1 mg/g to 12.5 mg/g in peel and 101.5 mg/g to 26.9 mg/g in pulp). In peel, glucose decreased by 79.10% in lightly infected fruits compared with healthy fruits (p < 0.0001) and remained 75.65% lower in moderately infected fruits. However, in highly infected peel, glucose was only 7.99% lower than in healthy fruits, with no significant difference (p > 0.05). In pulp, glucose decreased by 71.68% in lightly infected fruits (p < 0.0001) and by 24.92% in moderately infected fruits. In highly infected pulp, glucose was 6.9% higher than in healthy fruits; however, this increase was not statistically significant. Sucrose exhibited the greatest decline among the quantified sugars during the early stages of ASSD (36.3 mg/g to 3.8 mg/g in peel and 46.4 mg/g to 10.7 mg/g in pulp). In peel, sucrose decreased by 86.64% in lightly infected fruits compared with healthy fruits (p < 0.0001) and remained 40.58% lower in moderately infected fruits. However, sucrose levels in highly infected peel recovered to values comparable to those of healthy fruits, differing by only 0.04%. In pulp, sucrose decreased by 77.0% in lightly infected fruits (p < 0.0001) and remained 29.85% and 42.46% lower in moderately and highly infected fruits, respectively. The reduction in highly infected pulp remained statistically significant. Overall, the soluble sugars decreased sharply at the lightly infected stage in both tissues. Their subsequent patterns differed between peel and pulp: fructose, glucose, and sucrose returned to levels comparable to healthy samples in highly infected peel, whereas fructose and sucrose remained lower in highly infected pulp.

**Figure 3.**
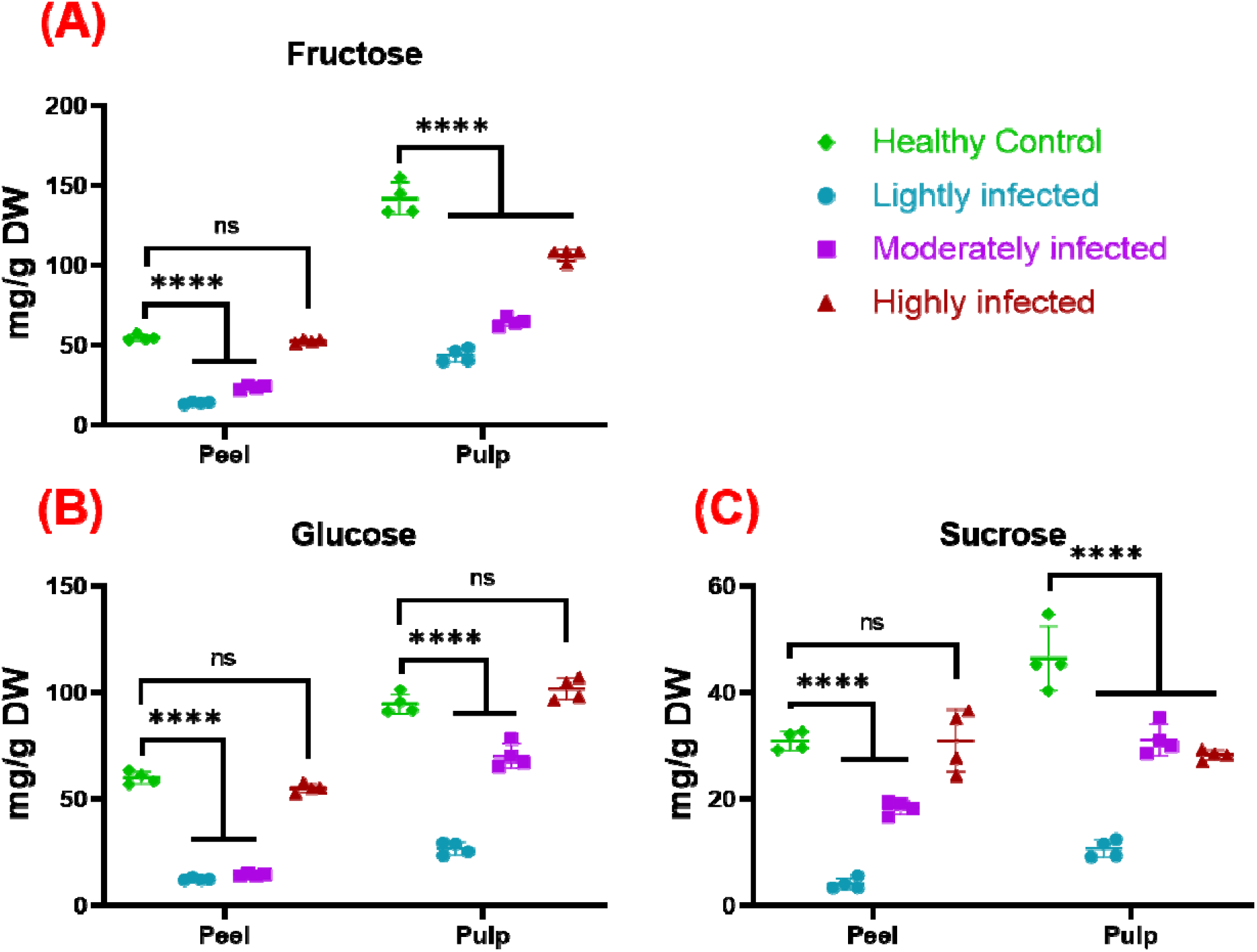
Concentrations of major soluble sugars in apple peel and pulp at different stage of apple scar skin disease (ASSD). Concentrations of (A) fructose, (B) glucose, and (C) sucrose were determined by ^1^H-NMR spectroscopy and are expressed as mg g^-1^ DW. (*n* = 4 biological replicates per stage).

The comparison of TSS with individual sugars showed different trends (Figure. 4). TSS increased at the moderate and high infection stages, whereas glucose, fructose, and sucrose decreased sharply at the lightly infected stage and subsequently increased to varying degrees. The same patterns were evident in the Z-score normalized data and fold changes relative to healthy fruits.

**Figure 4.**
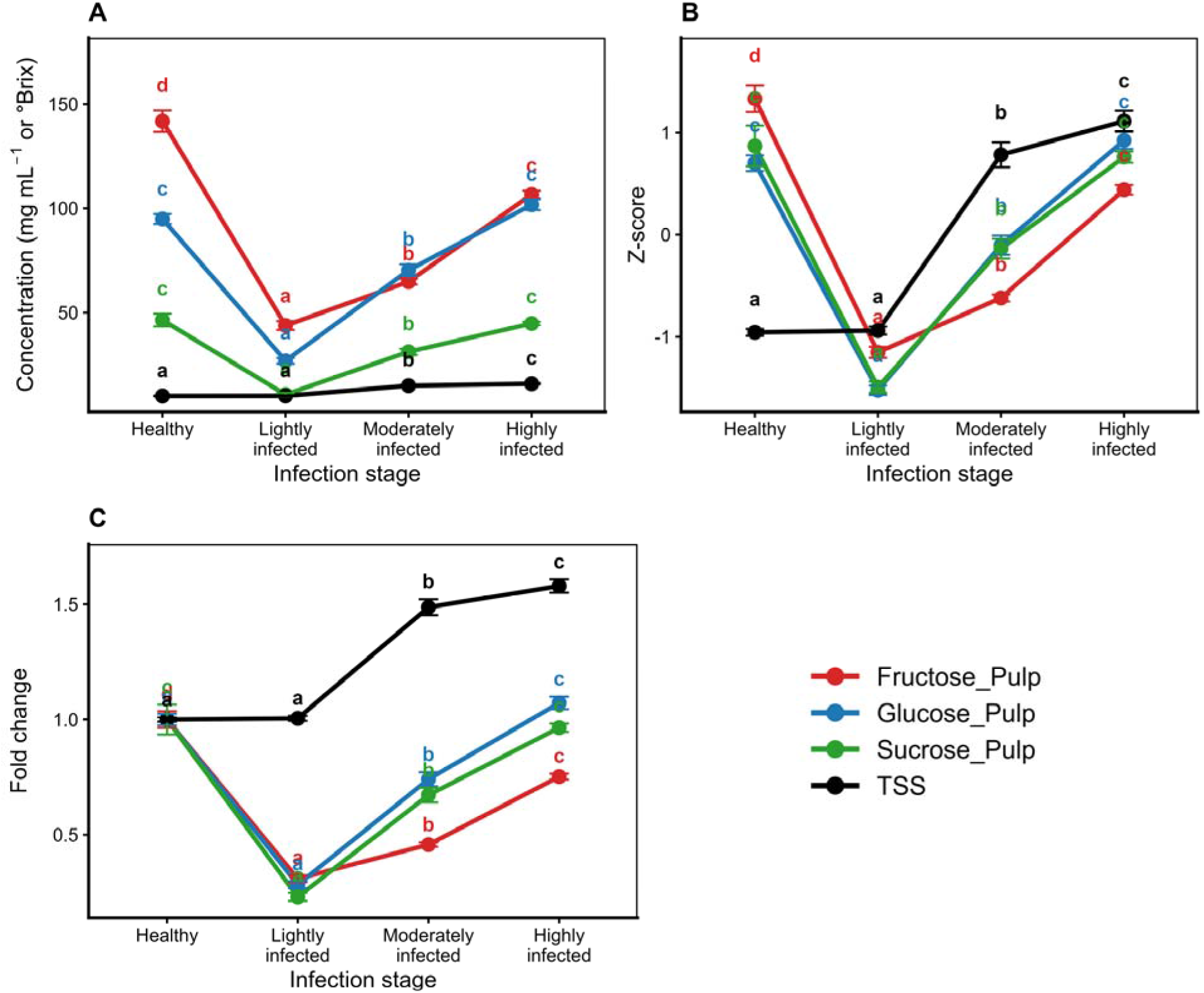
Comparison of total soluble solids and major soluble sugars during progression of apple scar skin disease (ASSD). **(A)** Absolute values of total soluble solids (TSS; °Brix) and glucose, fructose, and sucrose concentrations across healthy, lightly infected, moderately infected, and highly infected fruits. TSS was measured by refractometry (*n* = 10 biological replicates per stage), whereas individual sugars were quantified in pulp by ^1^H-NMR spectroscopy (*n* = 4 biological replicates per stage). **(B)** Z-score normalized values and **(C)** fold changes relative to the healthy group.

### 3.5 Multivariate analysis of GC-MS profiles

Representative overlaid GC-MS total ion chromatograms (TICs) of healthy control and ASSD-infected apple peel and pulp samples, including the complete retention time range of the identified metabolites, are presented in Supplementary Figures S4 and S5, respectively. Multivariate analyses of GC-MS profiles revealed clear, stage-dependent metabolic restructuring in both tissues (Figure. 5). PCA showed separation among infection stages in both peel and pulp (Figure. 5A, B). In peel, PC1 (45.8%) and PC2 (27.9%) together explained 73.7% of the total variance, whereas in pulp, PC1 (56.4%) and PC2 (15.8%) together accounted for 72.2%. Separation among infection stages was more apparent in peel, whereas highly infected pulp samples were positioned closer to the healthy group. PLS-DA showed separation among infection stages in both tissues (Figure. 5C, D). The peel model showed high explained and predicted variance (R² = 0.97, Q² = 0.95) and was supported by permutation testing (p < 0.01). The pulp model also showed high explained and predicted variance (R² = 0.93, Q² = 0.86); however, permutation testing was not significant (p = 0.4). The supervised separation observed for pulp should therefore be interpreted cautiously. VIP analysis identified the metabolites contributing most strongly to PLS-DA separation (Figure. 5E, F). In peel, metabolites with VIP scores > 1 included triterpenoids, phenolic-related compounds, lipids, sugars, and organic acids. In pulp, metabolites with VIP scores > 1 included amino acids, polyols, sugars, and organic acids.

**Figure 5.**
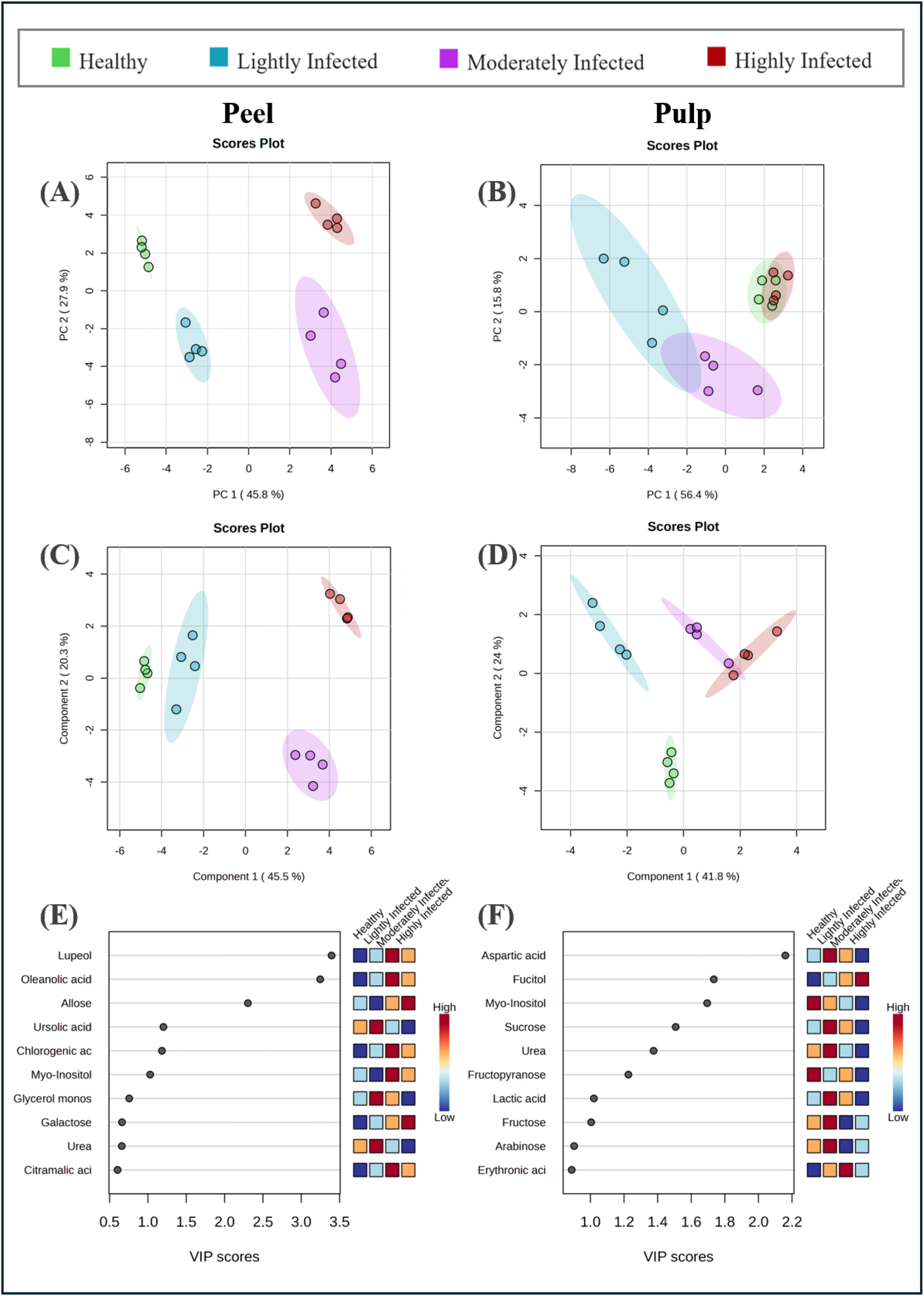
Multivariate analysis of GC-MS metabolomic profiles of apple peel and pulp across progressive stages of apple scar skin disease (ASSD). **(A, B)** Principal component analysis (PCA) score plots of peel and pulp samples, respectively, showing the distribution of healthy, lightly infected, moderately infected, and highly infected samples. Metabolite data were first normalized to the internal standard (ribitol), then subjected to median normalization, log□ transformation, and pareto scaling prior to multivariate analysis. The variance explained by each principal component (%) is indicated on the corresponding axis. **(C, D)** Partial least squares-discriminant analysis (PLS-DA) score plots of peel and pulp samples, respectively. The peel model showed high explained and predicted variance (R² = 0.97, Q² = 0.95) and was supported by permutation testing with 100 permutations (*p* < 0.01). The pulp model also showed high explained and predicted variance (R² = 0.93, Q² = 0.86); however, permutation testing was not significant (*p* = 0.4), and the observed supervised separation should therefore be interpreted with caution. **(E, F)** Variable importance in projection (VIP) scores for peel and pulp, respectively, showing the metabolites contributing most strongly to PLS-DA separation. Metabolites with VIP scores > 1 were considered important contributors to group separation.

### 3.6 Tissue-specific metabolic pathway alterations during ASSD progression

One-way ANOVA followed by FDR correction identified 25 and 21 metabolites that differed significantly among infection stages in peel and pulp, respectively (Supplementary Tables S1 and S2). In peel, metabolites showed distinct patterns across disease stages (Figure. 6). Gluconic acid, galactose, mannose, mannitol, and glycerol generally increased with increasing symptom severity, whereas fructose, fructopyranose, and fructofuranose were most abundant at the highly infected stage. Arabinose and sucrose decreased at the later infection stages. Glucose, glucopyranose, ribofuranose, sorbitol, allose, fucitol, and lactitol showed their highest relative abundance at the moderately infected stage. Among organic acids, citrate increased toward the later infection stages, whereas malate, quinic acid, and citramalate were highest at the moderately infected stage. Lactate was higher at the lightly and moderately infected stages but decreased at the highly infected stage. Chlorogenic acid and lupeol were most abundant in highly infected peel, whereas ursolic and oleanolic acids decreased with increasing disease severity. Stearic acid increased toward the highly infected stage, while palmitic acid was highest at the moderately infected stage and monopalmitin accumulated mainly at the highly infected stage.

**Figure 6.**
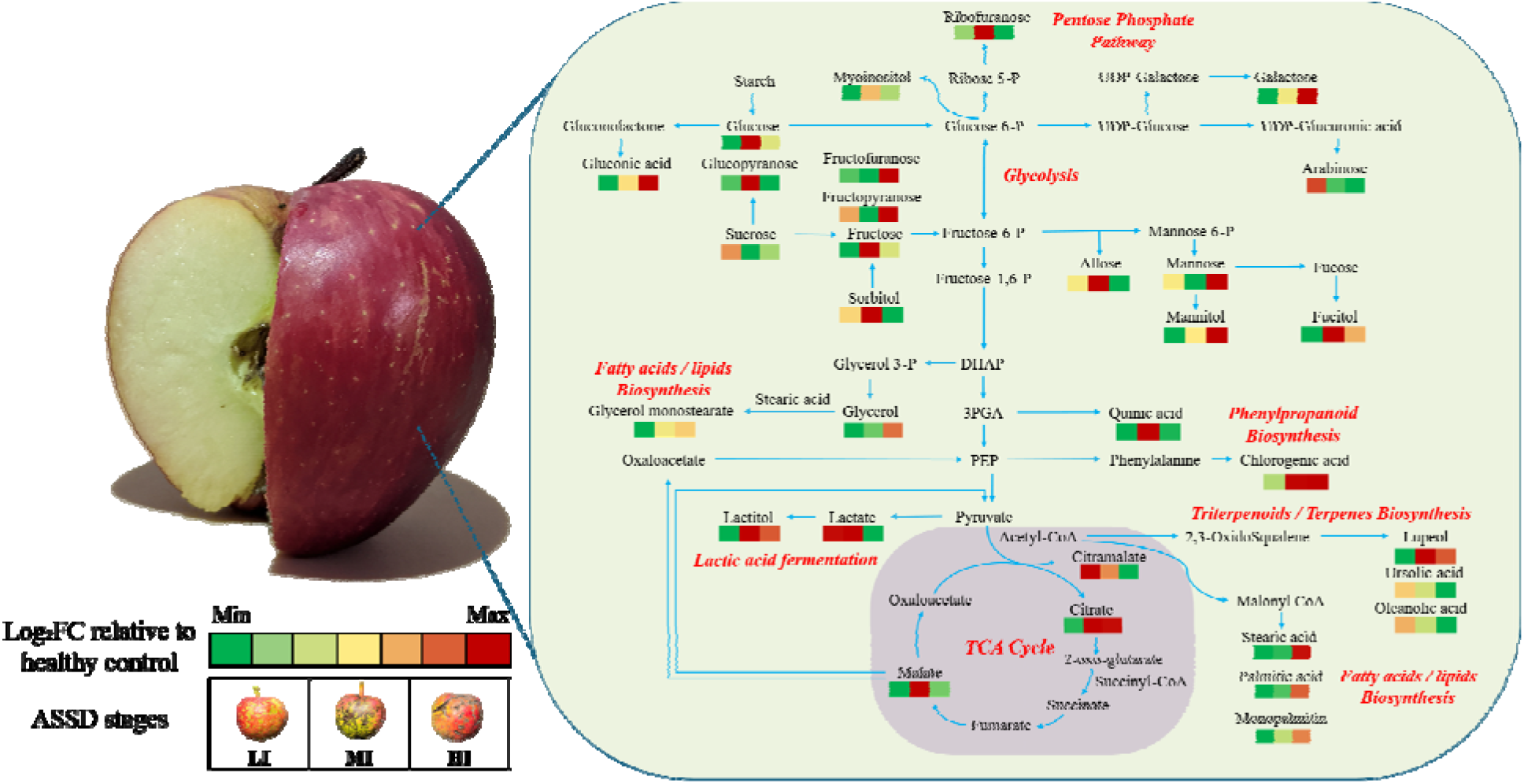
GC-MS based metabolic pathway map of apple peel during progression of apple scar skin disease (ASSD). Metabolite perturbations in lightly infected (LI), moderately infected (MI), and highly infected (HI) samples are expressed as log□ fold changes (log□FC) relative to the healthy group. For each metabolite, the three boxes represent LI, MI, and HI, respectively. Negative and positive values indicate lower and higher relative abundance, respectively, compared with healthy peel.

In pulp, several sugars and polyols also differed across disease stages (Figure. 7). Glucose, arabinose, sucrose, fructose, and fructopyranose generally decreased toward the highly infected stage, whereas glycerol increased with disease severity. Ribofuranose, sorbitol, and fucitol were most abundant at the moderately infected stage. Mannitol was higher at the lightly and moderately infected stages before decreasing at the highly infected stage, while myo-inositol and fructofuranose remained lower than in healthy pulp across the infected groups. Citrate, quinic acid, erythronic acid, and malate were most abundant at the moderately infected stage and subsequently decreased at the highly infected stage. Lactate decreased with increasing disease severity. Threonine and aspartate also decreased toward the highly infected stage. Chlorogenic acid was most abundant at the lightly infected stage and declined thereafter. Stearic acid increased toward the moderate and high infection stages, whereas monopalmitin increased progressively across the infected groups.

**Figure 7.**
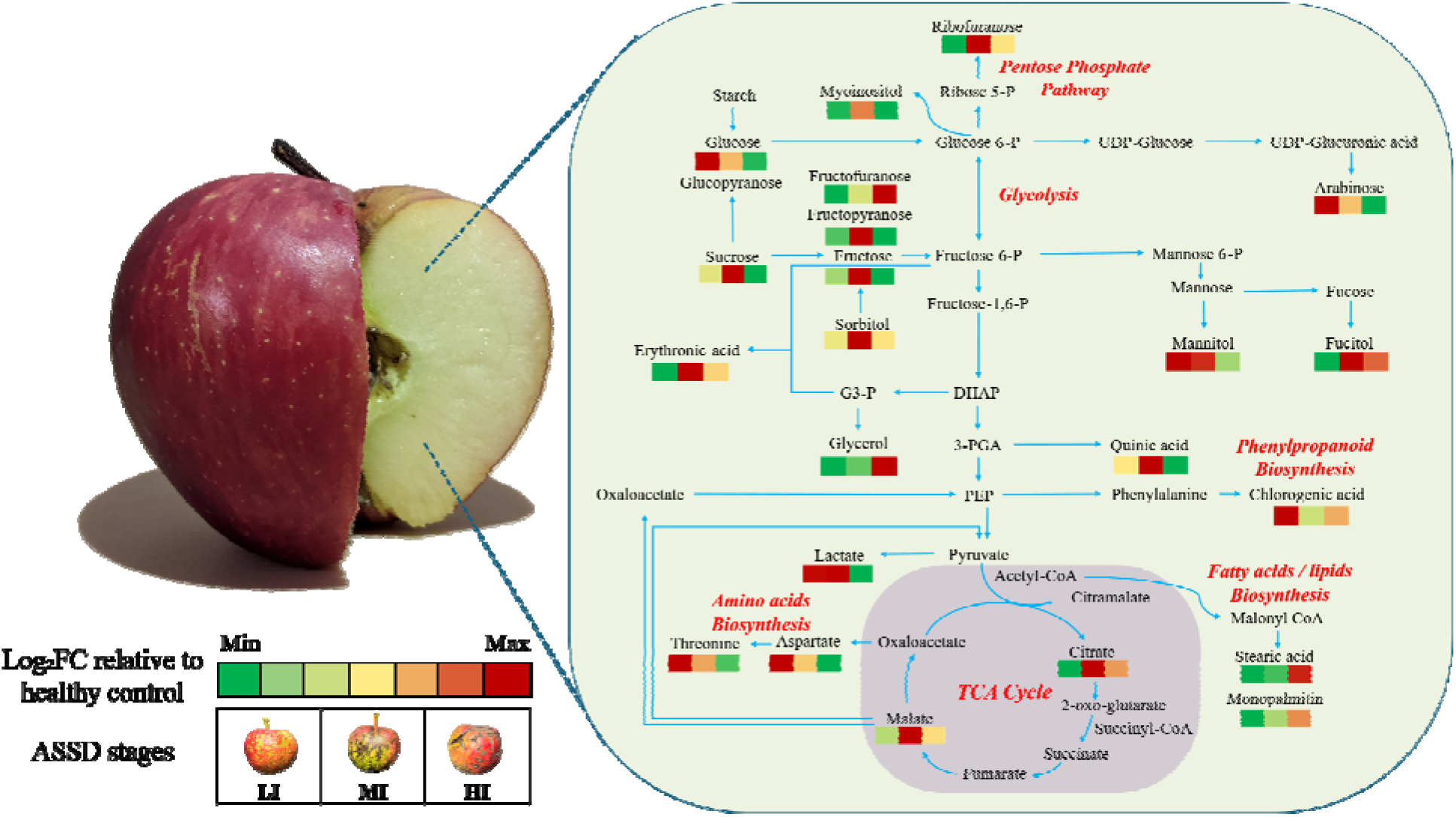
GC-MS based metabolic pathway map of apple pulp during progression of appl scar skin disease (ASSD). Metabolite alterations in lightly infected (LI), moderately infected (MI), and highly infected (HI) samples are expressed as log□ fold changes (log□ FC) relative to the healthy group. For each metabolite, the three boxes represent LI, MI, and HI, respectively. Negative and positive values indicate lower and higher relative abundance, respectively, compared with healthy peel.

### 3.7 Identification of candidate discriminatory metabolites

ROC analysis identified several metabolites that distinguished healthy from infected sample (Supplementary Figure. 3). In peel, the highest AUC values were obtained for lupeol (AUC = 1.00), sucrose (AUC = 0.979), ursolic acid (AUC = 0.958), and mannose (AUC = 0.958). In pulp, the corresponding metabolites were myo-inositol (AUC = 1.00), chlorogenic acid (AUC = 0.938), glucitol (sorbitol; AUC = 0.938), and monopalmitin (AUC = 0.896). Complete ROC analysis data for peel and pulp are available in Supplementary Tables S3 and S4, respectively. To identify the most informative candidate discriminatory metabolites, ROC performance was considered together with PLS-DA VIP scores and statistical significance across infection stages. Based on these combined criteria, lupeol, ursolic acid, and chlorogenic acid were selected in peel, whereas myo-inositol, chlorogenic acid, and aspartic acid were selected in pulp. These metabolites were therefore retained as the final candidate discriminatory metabolites for further evaluation.

## 4. Discussion

### 4.1 ASSVd sequence variation and symptom progression

Molecular analysis confirmed ASSVd infection in all symptomatic groups. The sequences obtained from lightly, moderately, and highly infected fruits clustered with different previously reported ASSVd isolates and occupied different positions in the phylogenetic analysis, indicating sequence variation among the isolates detected in the sampled orchard. However, because fruits were collected from naturally infected trees and classified according to symptom severity rather than obtained through controlled inoculation with defined ASSVd variants, the relationship between sequence variation and symptom severity could not be established. Consequently, the physicochemical and metabolic perturbations observed in this study are interpreted in relation to ASSD symptom progression rather than specific ASSVd sequence variants.

### 4.2 ASSD-induced perturbations in fruit quality and physicochemical characteristics

Apple scar skin disease progression was associated with changes in fruit morphology and physicochemical quality. The reductions in fruit weight, height, and firmness at the highly infected stage suggest that severe ASSD impairs normal fruit development, consistent with previous reports on viroid-infected apple fruit (Chen *et al*., 2023b; Hadidi *et al*., 2024; Li *et al*., 2023). The observed reduction in firmness may be associated with viroid-induced alterations in cellular organization and membrane integrity (Di Serio *et al*., 2013), although the underlying mechanism remains uncertain because cell-wall composition and membrane composition were not assessed in the present study. The progressive increase in titratable acidity suggests that ASSD is associated with changes in organic acid metabolism linked to central carbon metabolism, indicating that disease progression extends beyond visible symptoms on the fruit surface to influence overall fruit quality and composition (Chen *et al*., 2023a; Sharma *et al*., 2020).

### 4.3 Tissue-specific metabolic responses to ASSD progression

Metabolomic profiling showed distinct responses of peel and pulp during ASSD progression. As presented in Figure. 5, PCA distinguished the different disease stages in both tissues, although the separation was more pronounced in the peel. This pattern is expected because ASSD symptoms develop primarily on the fruit surface (He *et al*., 2026). Similar tissue-specific metabolic responses in apple peel have been documented in metabolomic studies of other apple diseases. (Falginella *et al*., 2021; Shah *et al*., 2023; He *et al*., 2026). In the present study, the major differences between tissues were observed for pentacyclic triterpenoids in the peel and for sugar alcohols and amino acids metabolism in the pulp, supports previous reports describing tissue-specific distributions of these metabolites (Falginella *et al*., 2021; Li *et al*., 2023; He *et al*., 2026). Together, these results indicate that ASSD affects the metabolism of peel and pulp differently, reflecting the distinct physiological functions of the two tissues.

### 4.4 ASSD-induced reorganization of carbohydrate metabolism

The disruption in soluble sugars during the early stages of ASSD indicate that carbohydrate metabolism is affected soon after infection. Rather than declining progressively with symptom severity, fructose, glucose, and sucrose reached their lowest levels in lightly infected fruit before showing partial, tissue-specific recovery at later stages. This pattern points to a dynamic adjustment of carbohydrate metabolism as the disease develops. A similar early depletion of soluble sugars has been reported by Hu *et al*. (2019) during latent *Botrytis cinerea* infection in strawberry, where it was attributed to increased carbohydrate consumption associated with defence responses. Soluble sugars function not only as carbon sources but also as signalling molecules that regulate plant defence and stress responses (Morkunas & Ratajczak, 2014). In addition, pathogen infection can alter sugar partitioning through the activity of sugar transporters and invertases, thereby modifying carbon allocation within host tissues (Liu *et al*., 2022). The partial recovery of sugar levels at later stages may therefore result from changes in carbohydrate transport, utilization, or redistribution as infection progresses (Rubio *et al*., 2021).

Although total soluble solids increased in moderately and highly infected fruit, the concentrations of the major soluble sugars did not show a corresponding increase. This discrepancy suggests that other soluble constituents, including sugar alcohols, organic acids, and amino acids etc, contributed to the elevated °Brix values (Nawaz *et al*., 2024). Consistent with the results, GC-MS analysis also revealed changes in sugar alcohol metabolism, particularly sorbitol. As the primary translocated carbohydrate in apple, sorbitol is central to source-sink carbon transport (Loescher *et al*., 1982), and the variation observed during ASSD is consistent with altered carbon partitioning in infected fruit.

### 4.5 Viroid-induced alterations in cuticular triterpenoid metabolism during ASSD progression

Among the peel-associated metabolites, pentacyclic triterpenoids exhibited distinct stage-specific responses during ASSD progression, indicating that triterpenoid metabolism is altered during disease development. Apple peel is naturally enriched in pentacyclic triterpenoids, which are major constituents of the cuticular wax and play essential roles in maintaining the structural integrity and protective functions of the fruit surface (Butkevičiūtė et al., 2022). In the present study, lupeol increased progressively with disease severity, whereas ursolic acid and oleanolic acid declined, suggesting differential modulation of individual branches of triterpenoid metabolism during ASSD progression. These contrasting responses likely reflect disease-associated metabolic reprogramming that influences cuticle maintenance and stress adaptation.

**Figure 9.**
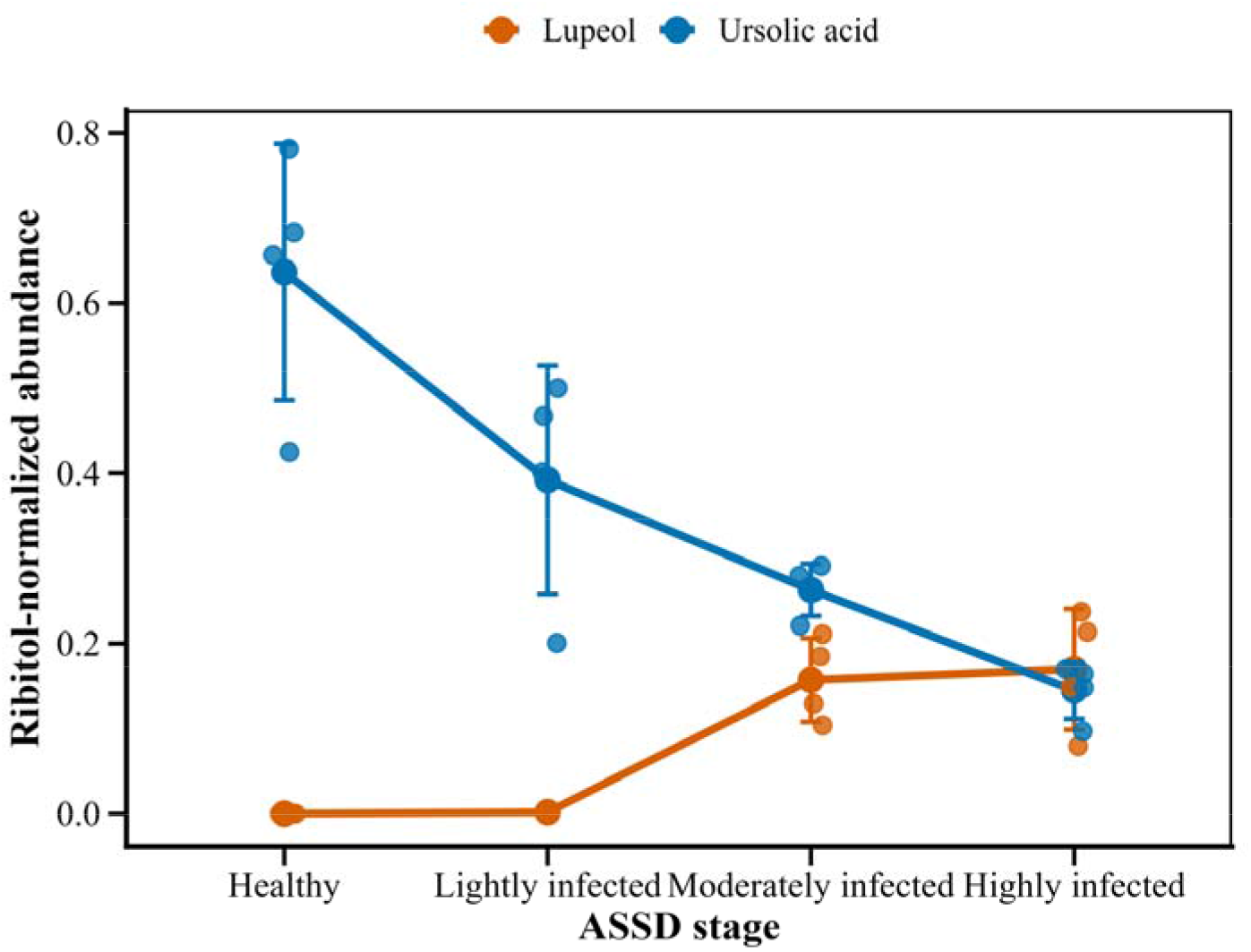
Relative abundances of the pentacyclic triterpenoids lupeol and ursolic acid during Apple Scar Skin Disease (ASSD) progression, illustrating the accumulation of lupeol and the depletion of ursolic acid with increasing disease severity.

Lupeol is synthesized through the mevalonate pathway and serves as an important intermediate in pentacyclic triterpenoid biosynthesis. Consequently, its increased abundance may reflect alterations in triterpenoid biosynthetic flux under pathogen-induced stress (Andre et al., 2016; Falginella et al., 2021). Although lupeol is a minor constituent of apple cuticular wax compared with ursolic and oleanolic acids, it has been reported to possess antimicrobial, antifungal, and antioxidant activities, and its accumulation has been associated with pathogen infection, oxidative stress, and wound responses in several plant species (Shu et al., 2019; Javed et al., 2021). Therefore, the progressive accumulation of lupeol observed during ASSD progression may represent an adaptive metabolic response associated with enhanced defence-related metabolism. In contrast, ursolic acid and oleanolic acid declined progressively with increasing disease severity. These pentacyclic triterpenoids are the predominant constituents of apple cuticular wax and contribute substantially to cuticle architecture, mechanical stability, hydrophobicity, and permeability, thereby minimizing transpirational water loss and preserving postharvest fruit quality (Dashbaldan et al., 2020; Butkevičiūtė et al., 2022). As the cuticle constitutes the primary physical barrier against pathogen invasion, reductions in these triterpenoids may reflect disease-associated alterations in cuticular wax composition that could compromise the protective properties of the fruit surface (Falginella et al., 2021). Ursolic acid has additionally been implicated in maintaining cuticular stability under oxidative stress, desiccation, and ultraviolet radiation (Zhang et al., 2020), whereas oleanolic acid contributes to wax organization, cuticle permeability, and is widely recognized as a biochemical marker of peel triterpenoid metabolism (Zhang et al., 2020). The simultaneous decline of ursolic and oleanolic acids together with the accumulation of lupeol therefore suggests a redistribution of triterpenoid metabolism during ASSD progression, potentially reflecting disease-associated remodeling of cuticular wax composition.

Viroids are known to extensively reprogram host gene expression and secondary metabolic pathways through RNA-mediated regulatory mechanisms rather than by directly degrading metabolites (Moyo et al., 2022). Therefore, the progressive depletion of ursolic acid observed during ASSD progression is more likely to result from viroid-induced modulation of host triterpenoid biosynthesis and cuticular wax metabolism than from direct metabolite degradation. Such metabolic reprogramming may alter the accumulation of cuticle-associated triterpenoids, thereby contributing to the reduced structural integrity and barrier function of the fruit surface and facilitating the development of characteristic ASSD symptoms, including scarring and cracking. This interpretation is consistent with previous studies demonstrating that alterations in triterpenoid biosynthesis influence apple cuticle composition and integrity during fruit development and skin disorders (Falginella et al., 2021). Nevertheless, because neither cuticle ultrastructure, wax composition, nor the expression of triterpenoid biosynthetic genes was examined in the present study, the observed metabolic changes should be interpreted as evidence of altered host triterpenoid metabolism rather than conclusive evidence of impaired cuticle structure or function.

### 4.6 Tissue-specific phenolic and phenylpropanoid responses to ASSD

The peel exhibited greater changes in phenolic metabolism than the pulp, consistent with its role as the principal site of phenolic accumulation in apple fruit. The transient increase in total phenolic content during the early stages of ASSD, followed by a decline in highly infected fruit, suggest that phenylpropanoid metabolism is initially stimulated but becomes less active as disease severity increases. Comparable biphasic responses have also been described in other apple fruit disorders. During the development of superficial scald, Cebulj *et al*. (2020) observed an early induction of phenylpropanoid-related genes, whereas the abundance of most polyphenol groups and the activities of flavonoid biosynthetic enzymes declined as symptoms became more severe, likely because of increased phenolic turnover. A similar pattern was reported by Schovánková *et al*. (2011), who found that fungal infection initially stimulated phenolic accumulation and phenylalanine ammonia-lyase (PAL) activity, followed by a gradual decline as disease progressed. Together, these observations suggest that phenylpropanoid metabolism in apple fruit is dynamic, with an early activation of phenolic biosynthesis that is subsequently followed by reduced biosynthetic activity and/or enhanced phenolic turnover during disease development. The tissue-specific behaviour of chlorogenic acid in the present study is consistent with this pattern. Chlorogenic acid, a major phenylpropanoid metabolite with antioxidant and antimicrobial functions (Jiao *et al*., 2018; Romanazzi *et al*., 2016; Ilea *et al*., 2026), accumulated during the early stages of infection in the pulp but increased progressively in the peel. This difference suggests that phenylpropanoid metabolism is regulated differently between tissues and that defence-related metabolic activity may be sustained for longer in the peel than in the pulp. These observations are consistent with the transcriptomic findings of He *et al*. (2026), who demonstrated that ASSVd perturbs phenylpropanoid metabolism by suppressing genes involved in anthocyanin biosynthesis, supporting differential regulation of phenylpropanoid metabolism between peel and pulp during ASSD progression.

### 4.7 Metabolic alterations in aspartate-family amino acid metabolism and carbon-nitrogen metabolism

The concurrent decline in aspartate and threonine toward the highly infected stage in pulp suggests perturbation of aspartate-family amino acid metabolism during ASSD progression. Aspartate occupies a central position in carbon and nitrogen metabolism and, through its close metabolic connection with the tricarboxylic acid (TCA) cycle, serves as the precursor for the aspartate-family amino acids, including threonine, methionine, lysine, and isoleucine (Kirma *et al*., 2012; de la Torre *et al*., 2014; Li *et al*., 2017; Famiani *et al*., 2020). Therefore, the concurrent alterations in aspartate, threonine, and TCA cycle-associated organic acids indicate coordinated metabolic adjustments in carbon and nitrogen metabolism during ASSD progression.

### 4.8 Coordinated metabolic shifts in lipid and central carbon metabolism

The coordinated perturbations in fatty acids and glycerolipids suggest lipid metabolism is altered during ASSD progression. The altered abundance of palmitic acid, stearic acid, and monopalmitin may reflect changes in membrane lipid metabolism during ASSD progression. Similar responses have been reported in other plant-pathogen interactions, where membrane lipid remodelling contributes to stress signalling and maintenance of membrane integrity (Fan *et al*., 2018; Siebers *et al*., 2016; Li *et al*., 2024).

In central carbon metabolism, polyols exhibited stage-dependent responses. The accumulation of mannitol at later infection stages is consistent with its proposed roles in osmotic adjustment and reactive oxygen species scavenging during biotic stress (Williamson *et al*., 2002). Likewise, the elevated galactose levels observed in symptomatic tissues may reflect increased turnover of galactan-rich pectic side chains during cell-wall remodelling (Gross *et al*., 1979; Yoshioka *et al*., 1995), a process reported to accompany both ripening-associated softening and stress-induced metabolic reprogramming in apple fruit (Li *et al*., 2023; Reckleben *et al*., 2025). Elevated gluconic acid may similarly reflect enhanced carbohydrate oxidation under stress conditions (Baxter *et al*., 2007; Silva *et al*., 2020). Together, these observations are consistent with previous reports indicating that perturbations in polyol and carbohydrate metabolism contribute to defence-related metabolic reprogramming during plant-pathogen interactions (Kanayama, 2009; Meng *et al*., 2018; Morkunas *et al*., 2014; Tauzin & Giardina 2014).

The stage-dependent perturbations in organic acids indicate that ASSD affects central carbon metabolism during disease progression. The accumulation of citrate at later infection stages, together with alterations in malate and lactate, is consistent with perturbations in the tricarboxylic acid (TCA) cycle and associated respiratory metabolism (Shah *et al*., 2023; Žebeljan *et al*., 2021). The **c**orresponding increase in titratable acidity further supports disruption of organic acid homeostasis. The transient accumulation of lactate during the early stages of infection may reflect shifts toward glycolytic or fermentative metabolism under stress conditions (Jiang *et al*., 2023). Moreover, intermediates of the tricarboxylic acid (TCA) cycle serve as carbon skeletons for amino acid biosynthesis through enzymes such as aspartate aminotransferase (Famiani *et al*., 2020); the observed perturbations in organic acids are consistent with the concurrent perturbation in amino acid metabolism, indicating coordinated reprogramming of carbon and nitrogen metabolism during ASSD progression.

### 4.9 Candidate discriminatory metabolites

The candidate discriminatory metabolites identified in this study represent distinct metabolic processes associated with ASSD progression. Peel candidates (lupeol, ursolic acid, and chlorogenic acid) reflect perturbations in triterpenoid and phenylpropanoid metabolism. Lupeol has been associated with antimicrobial and protective activities, whereas ursolic acid is a major constituent of the apple cuticular wax, and chlorogenic acid is a key phenylpropanoid-derived metabolite with antioxidant and antimicrobial properties (Shu *et al*., 2019; Javed *et al*., 2021; Dashbaldan *et al*., 2020; Jiao *et al*., 2018; Romanazzi *et al*., 2016; Ilea *et al*., 2026). In contrast, the pulp candidates (myo-inositol, chlorogenic acid, and aspartate) are associated with primary metabolism, including cellular signalling, osmotic regulation, cell-wall metabolism, and carbon-nitrogen metabolism (Loewus & Murthy, 2000; Endres & Tenhaken, 2009; de la Torre *et al*., 2014; Li *et al*., 2017; Famiani *et al*., 2020). The occurrence of chlorogenic acid in both tissues further suggests that phenylpropanoid metabolism is consistently altered during ASSD progression, although its contrasting accumulation patterns indicate tissue-specific regulation (He *et al*., 2026). Collectively, these metabolites represent promising candidate discriminatory metabolites associated with ASSD progression. However, their diagnostic utility should be validated in larger, independent sample sets encompassing multiple cultivars, orchards, and growing seasons before they can be considered reliable biomarkers.

## 5. Conclusion

Apple scar skin disease (ASSD) was associated with progressive alterations in fruit morphology, physicochemical characteristics, and metabolism, with the amplitude of these changes increasing with disease severity. Highly infected fruits exhibited reduced fruit weight, height, and firmness, accompanied by increased titratable acidity, indicating a progressive deterioration of fruit quality. Metabolomic analyses further revealed distinct tissue-specific responses to ASSVd infection, with the peel exhibiting more pronounced alterations in pentacyclic triterpenoids and phenolic compounds, whereas the pulp showed greater perturbations in sugars, polyols, amino acids, and organic acids. These findings demonstrate that ASSVd infection induces extensive metabolic reprogramming that extends beyond the development of visible peel symptoms. A key finding of this study was the contrasting accumulation pattern of peel-associated pentacyclic triterpenoids, characterized by the progressive accumulation of lupeol and the depletion of ursolic acid and oleanolic acid during ASSD progression. These coordinated changes suggest that ASSVd infection is associated with alterations in host triterpenoid metabolism and cuticular wax composition, potentially affecting the structural integrity and protective functions of the fruit surface. Although the underlying molecular mechanisms remain to be elucidated, these results provide new insights into the metabolic basis of ASSD progression and identify triterpenoid metabolism as a promising target for future mechanistic investigations.

Overall, among all the metabolites identified, lupeol, ursolic acid, and chlorogenic acid in the peel, together with myo-inositol, chlorogenic acid, and aspartic acid in the pulp, consistently discriminated healthy and infected fruits, highlighting their potential as candidate metabolic markers of disease progression. To our knowledge, this study represents the first comprehensive integration of GC-MS metabolomics, ^1^H NMR spectroscopy, and physicochemical analyses to characterize tissue-specific metabolic alterations across successive stages of ASSD progression. Future studies integrating metabolomic, transcriptomic, and proteomic approaches with analyses of cuticular wax composition and ultrastructure, together with validation across diverse cultivars, orchards, and growing seasons, will be essential to elucidate the molecular mechanisms underlying ASSVd-induced metabolic reprogramming and to establish robust biomarkers for the early detection and monitoring of Apple Scar Skin Disease.

## Credit authorship contribution statement

Prem Chand (PC): performed metabolic and biochemical experiments, writing original draft and conceptualization. Santosh Watpade (SW) and Shyam Kumar Masakapalli (SKM): review and editing, conceptualization and supervision. Hema Kumar, Esha Devi, and Rishav Kumar: Methodology & Investigation.

## Data availability statement

Data will be made available on request.

## Acknowledgement

SKM acknowledges the Himachal Pradesh State Agricultural Marketing Board (HPSAMB), Himachal Pradesh, India, for financial support (Ref. No. IITM/HPSAMB/HT/326). PC thanks the Ministry of Education (MoE), Government of India, and the Indian Institute of Technology Mandi (IIT Mandi), India, for the doctoral research fellowship. PC and SKM are thankful to BioX Centre and Advanced Materials Research Centre (AMRC), IIT Mandi, for providing access for research facilities. SW acknowledges the ICAR-Indian Agricultural Research Institute, Regional Station, Shimla, India, for providing the research facilities necessary to carry out this study.

