## Supplementary file for "Apple scar skin viroid disease induced physicochemical and metabolic alterations in Apples"

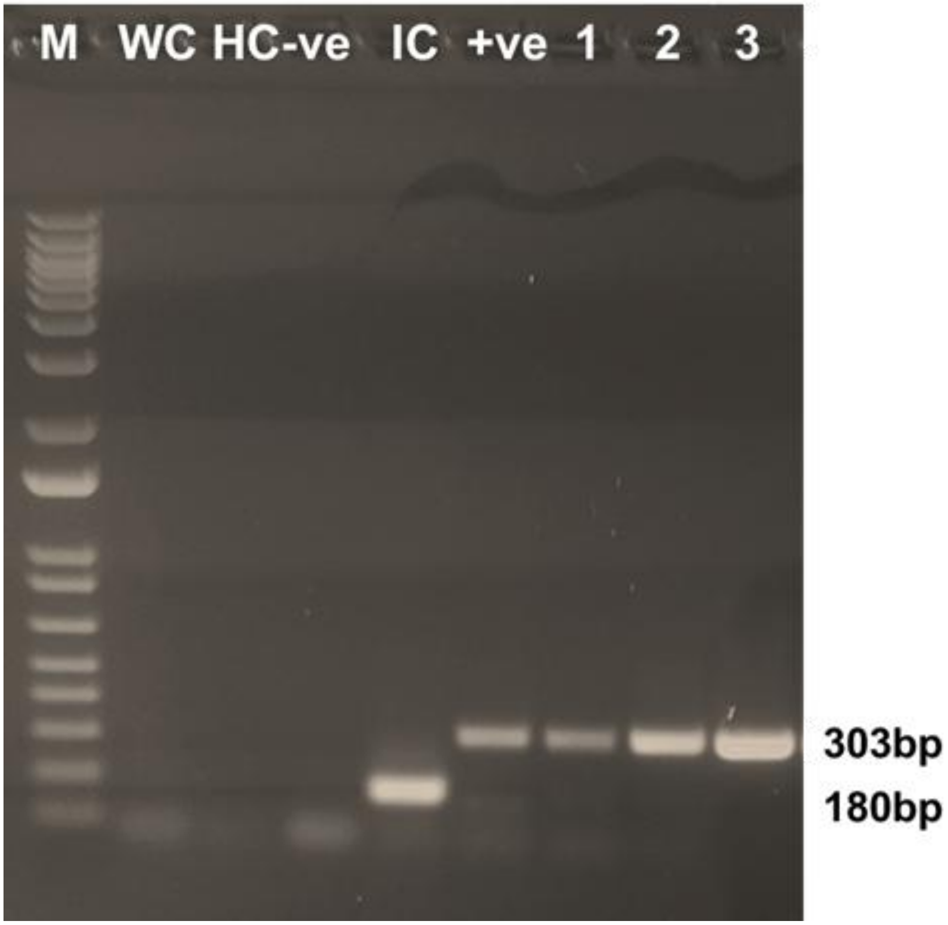


**Supplementary Figure 1**: RT-PCR amplification of viroid suspected apple fruit samples. Lane M: Marker, WC: water control, HC: healthy control, -ve: negative control, IC: internal control, +ve: positive control, 1: fruit sample LI, 2: fruit sample MI. 3: fruit sample HI

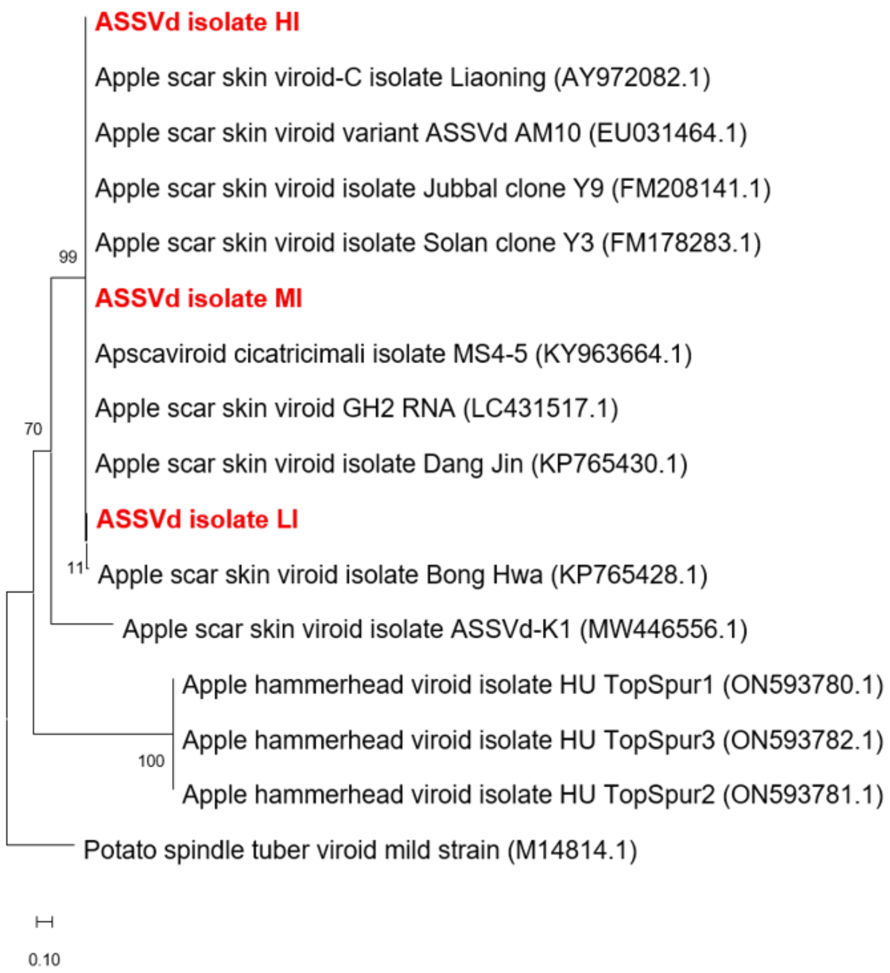


**Supplementary Figure 2**: Phylogenetic relationships among Apple scar skin viroid isolates were inferred using the maximum likelihood method. Branch support was assessed by bootstrap analysis with 1,000 replicates, and values are presented as percentages. Potato spindle tuber viroid was used as the outgroup.

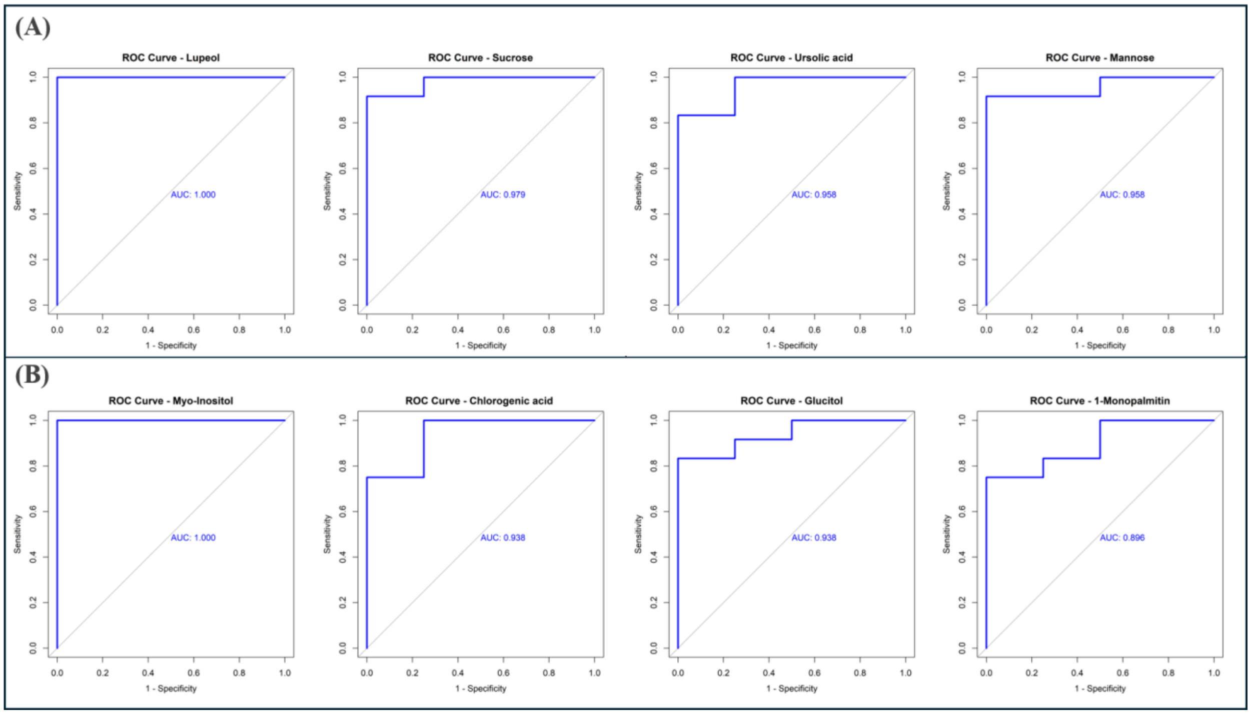


**Supplementary Figure 3:** ROC curves of candidate metabolites exhibiting the highest AUC values for discriminating healthy and ASSD-infected apple samples. ROC curves are shown for the four top-performing metabolites based on AUC values in **(A)** peel, and **(B)** pulp. The peel metabolites were lupeol, sucrose, ursolic acid, and mannose, whereas the pulp metabolites were myo-inositol, chlorogenic acid, glucitol (sorbitol), and monopalmitin. Discriminatory performance was assessed using the area under the receiver operating characteristic (ROC) curve (AUC), with 95% confidence intervals calculated by bootstrap resampling (2,000 iterations).

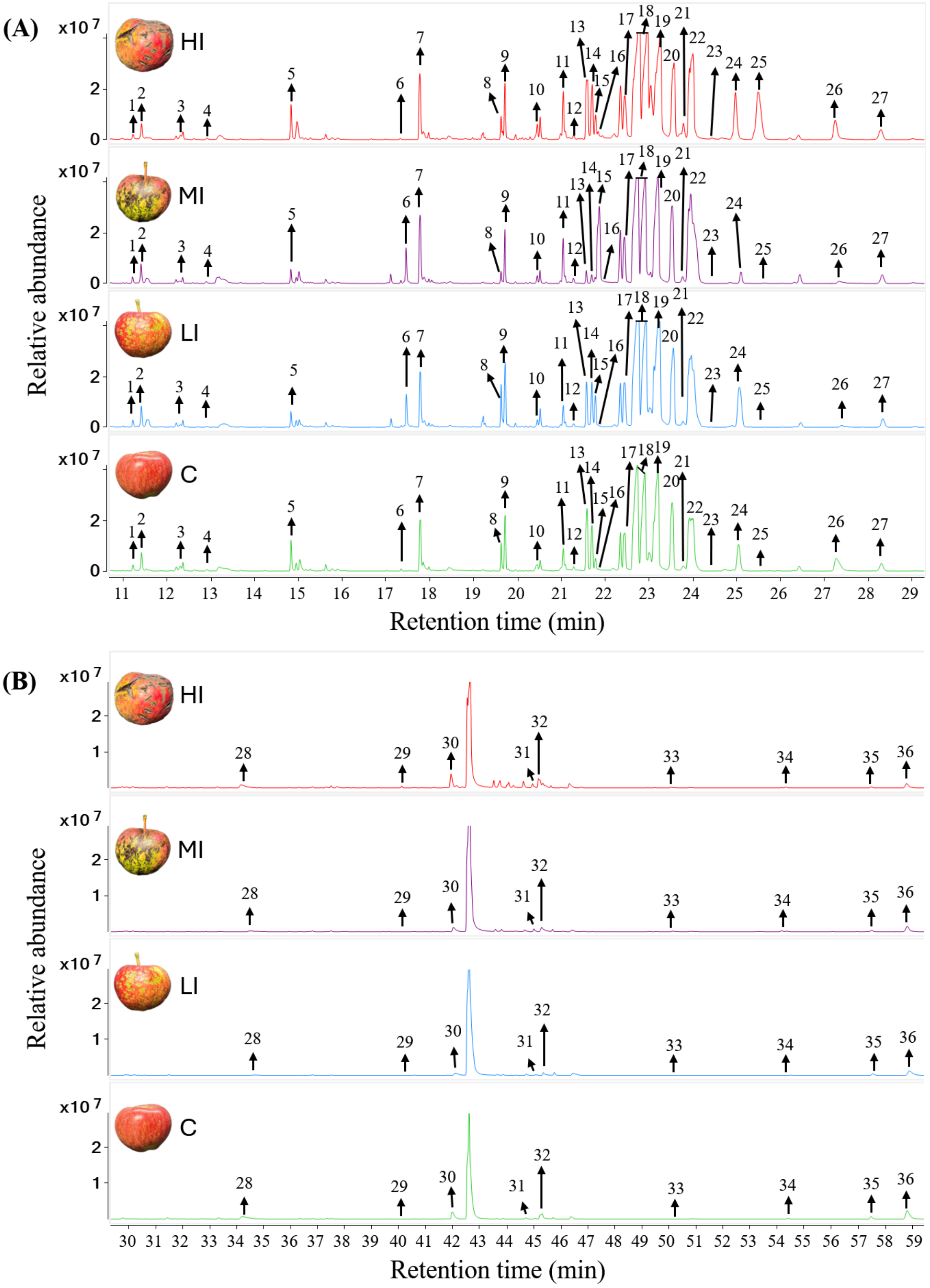


**Supplementary Figure S4.** Representative overlaid total ion chromatograms (TICs) of derivatized metabolites detected by GC-MS in healthy control and Apple Scar Skin Disease (ASSD) infected apple peel samples. Chromatograms are shown for the retention time intervals of

(A) 11–29 min and (B) 30–59 min, covering the complete range of identified metabolites. Ribitol was used as the internal standard (IS) for normalization. The metabolites identified correspond to the number as follows- 1:1,3-Propanediol, 2: Lactic acid, 3: Hydroxlyamine, 4: Urea, 5: Glycerol, 6: Citramalic acid, 7: Malic acid, 8: Arabinose, 9: Gluconic acid, 10: Ribitol (IS), 11: Fucitol, 12; Ribofuranose, 13 & 14 :Fructofuranose, 15 Fructopyranose, 16: Citric acid, 17: Quinic acid,18: Fructose, 19: Glucose, 20: Allose, 21: Mannitol, 22: Glucitol, 23: Mannose, 24: Glucopyranose, 25: Gluconic acid, 26: Palmitic Acid, 27: Myo-Inositol, 28: Stearic acid, 29: Sucrose, 30: 1-Monopalmitin, 31: Lactitol, 32: Glycerol monostearate, 33: Chlorogenic acid, 34: Lupeol, 35: Oleanolic acid, 36: Ursolic acid.

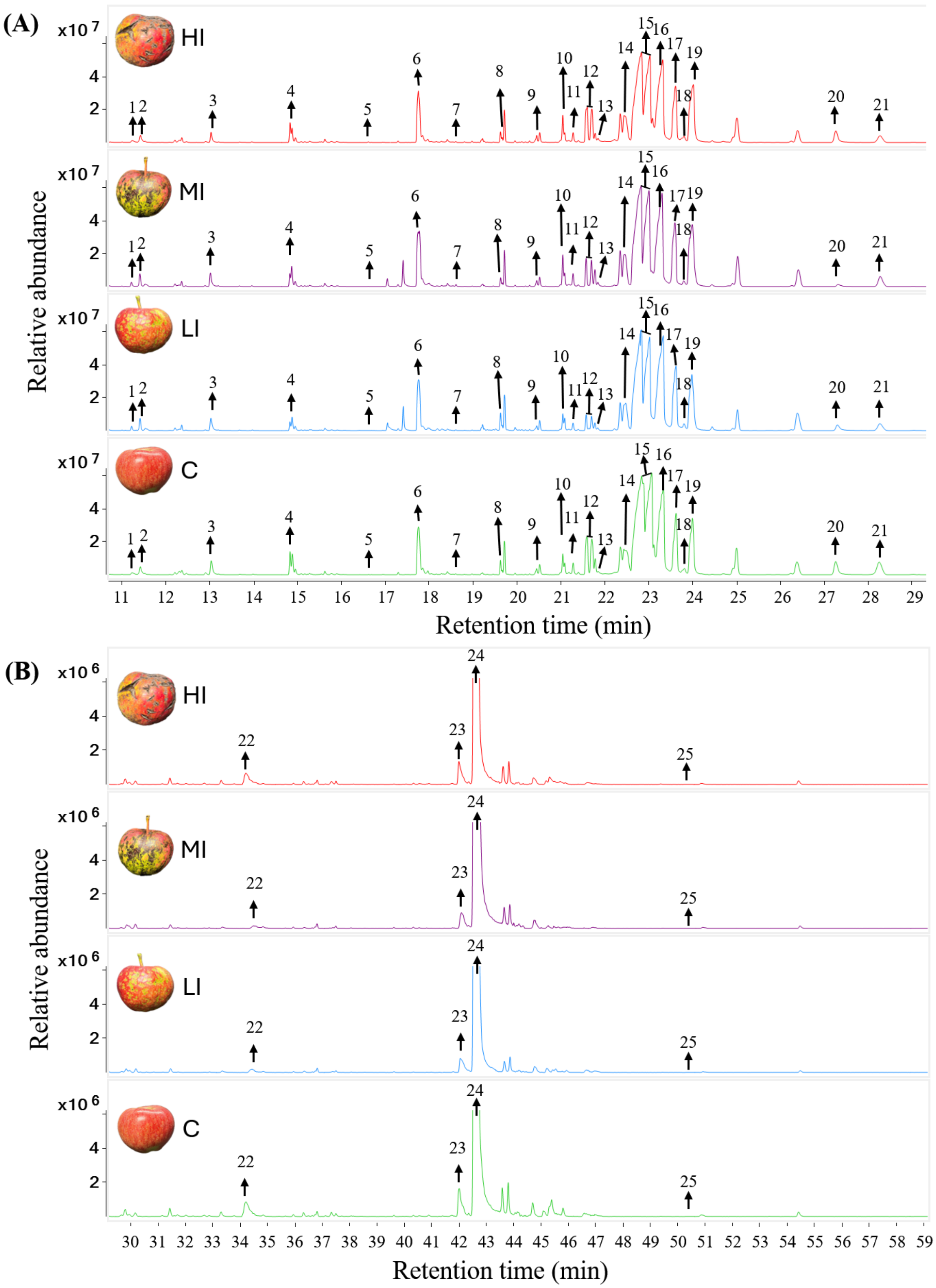


**Supplementary Figure S5**. Representative overlaid total ion chromatograms (TICs) of derivatized metabolites detected by GC-MS in healthy control and Apple Scar Skin Disease (ASSD) infected apple pulp samples. Chromatograms are shown for the retention time intervals of (A) 11–29 min and (B) 30–59 min, covering the complete range of identified metabolites. Ribitol was used as the internal standard (IS) for normalization. The metabolites identified correspond to the number as follows- 1: 1,3-Propanediol, 2: Lactic acid, 3: Urea, 4: Glycerol, 5: Threonine, 6: Malic acid, 7: Aspartic acid, 8: Arabinose, 9: Ribitol (IS), 10: Fucitol, 11; Ribofuranose, 12: Fructofuranose, 13: Citric acid, 14: Quinic acid,15: Fructose,16: Glucose, 17: Allose,18: Mannitol, 19: Glucitol, 20: Palmitic Acid, 21: Myo-Inositol, 22: Stearic acid, 23: 1-Monopalmitin, 24: Sucrose 25: Chlorogenic acid

**Supplementary Table S1**. Comprehensive summary of metabolite exhibiting statistically significant alterations (FDR < 0.05) across infection stages in apple peel tissue. The table includes stage-wise mean concentrations, $\log_2$ fold change values relative to healthy controls, p-values, and false discovery rate (FDR)-adjusted p-values for each metabolite.

| Metabolite | Mean Healthy | Mean Lightly | Mean Moderately | Mean Highly | Log2FC Lightly vs Healthy | Log2FC Moderately vs Healthy | Log2FC Highly vs Healthy | P value | FDR |
|---|---|---|---|---|---|---|---|---|---|
| Oleanolic acid | -16.7575 | -17.1177 | -2.68442 | -5.22942 | -0.36022 | 14.07308 | 11.52808 | 4.85E-17 | 1.65E-15 |
| Allose | -4.16029 | -5.92832 | -3.42461 | 3.284172 | -1.76803 | 0.735678 | 7.444459 | 1.10E-14 | 1.88E-13 |
| Lupeol | -16.7575 | -9.17428 | -2.72571 | -2.67719 | 7.583223 | 14.03179 | 14.08031 | 2.52E-14 | 2.86E-13 |
| Citramalic acid | 1.086346 | 0.350517 | 1.751334 | 1.353288 | -0.73583 | 0.664988 | 0.266942 | 1.82E-08 | 1.54E-07 |
| 1-Monopalmitin | 1.355376 | -0.03302 | 0.247699 | 1.010917 | -1.3884 | -1.10768 | -0.34446 | 1.19E-06 | 8.12E-06 |
| Glycerol monostearate | -5.00068 | -2.92563 | -3.79062 | -6.8187 | 2.075048 | 1.21006 | -1.81802 | 2.17E-06 | 1.05E-05 |
| Urea | 2.823041 | 2.695894 | 2.10034 | 1.976735 | -0.12715 | -0.7227 | -0.84631 | 2.13E-06 | 1.05E-05 |
| Chlorogenic acid | -2.70846 | -2.58916 | -1.07103 | -1.10626 | 0.119307 | 1.63743 | 1.602201 | 6.41E-06 | 2.72E-05 |
| Myo-Inositol | -2.91389 | -3.6986 | -1.51131 | -1.83706 | -0.78471 | 1.402585 | 1.076829 | 1.34E-05 | 5.05E-05 |
| Lactic acid | -3.03477 | -2.46766 | -3.46478 | -2.40241 | 0.56711 | -0.43001 | 0.632367 | 3.42E-05 | 0.000116 |
| Glucopyranose | 4.158072 | 4.33044 | 5.135124 | 4.058627 | 0.172368 | 0.977052 | -0.09945 | 5.56E-05 | 0.000172 |
| Ursolic acid | -0.68712 | -1.43333 | -1.93474 | -2.82223 | -0.74622 | -1.24763 | -2.13512 | 7.69E-05 | 0.000218 |
| Ribofuranose | 0.090144 | 0.111121 | 0.172728 | 0.466996 | 0.020976 | 0.082584 | 0.376852 | 0.0001 | 0.000262 |
| Glycerol | -1.40746 | -0.26117 | -0.2342 | -1.4715 | 1.146291 | 1.173259 | -0.06404 | 0.000261 | 0.000632 |
| Gluconic acid | 3.3476 | 1.950509 | 1.605578 | 3.465306 | -1.39709 | -1.74202 | 0.117707 | 0.000279 | 0.000632 |
| Stearic acid | 1.506837 | 0.653952 | 2.005458 | 0.179811 | -0.85289 | 0.498621 | -1.32703 | 0.000373 | 0.000792 |
| Sucrose | 1.210058 | 0.442385 | 0.83842 | 0.650503 | -0.76767 | -0.37164 | -0.55956 | 0.000424 | 0.000848 |

| | | | | | | | | | |
|---|---|---|---|---|---|---|---|---|---|
| Mannose | 2.419132 | -0.00993 | 0.338374 | 1.891494 | -2.42907 | -2.08076 | -0.52764 | 0.000483 | 0.000913 |
| Hydroxylamine | 2.545441 | 2.17918 | 2.859449 | 2.287382 | -0.36626 | 0.314008 | -0.25806 | 0.00112 | 0.002005 |
| 1,3-Propanediol | 0.680808 | 0.417881 | 0.864005 | 0.194687 | -0.26293 | 0.183197 | -0.48612 | 0.012777 | 0.021721 |
| Fucitol | 0.530204 | -2.06885 | -1.83696 | 0.449878 | -2.59906 | -2.36716 | -0.08033 | 0.014501 | 0.023479 |
| Fructose | 3.562604 | 3.546746 | 3.784939 | 3.316754 | -0.01586 | 0.222335 | -0.24585 | 0.020661 | 0.031479 |
| Galactose | 0.0031 | -0.27395 | 0.018755 | 0.630761 | -0.27705 | 0.015655 | 0.62766 | 0.021295 | 0.031479 |
| Quinic acid | 2.534768 | 2.297985 | 2.617476 | 2.310541 | -0.23678 | 0.082709 | -0.22423 | 0.031549 | 0.044695 |

**Supplementary Table S2.** Comprehensive summary of metabolite exhibiting statistically significant alterations (FDR < 0.05) across infection stages in apple pulp tissue. The table includes stage-wise mean concentrations, $\log_2$ fold change values relative to healthy controls, p-values, and false discovery rate (FDR)-adjusted p-values for each metabolite.

| Metabolite | Mean Healthy | Mean Lightly | Mean Moderately | Mean Highly | $Log_2FC$ Lightly vs Healthy | $Log_2FC$ Moderately vs Healthy | $Log_2FC$ Highly vs Healthy | P value | FDR |
|---|---|---|---|---|---|---|---|---|---|
| Fucitol | 0.859018 | 0.710308 | 1.769729 | 1.520413 | -0.14871 | 0.910711 | 0.661395 | 3.63E-10 | 8.71E-09 |
| Erythronic acid | -3.50769 | -3.55386 | -2.17196 | -3.2532 | -0.04617 | 1.335735 | 0.254492 | 1.39E-07 | 1.67E-06 |
| Threonine | -6.81448 | -4.60471 | -5.68024 | -6.60619 | 2.20977 | 1.134241 | 0.208293 | 1.13E-06 | 9.05E-06 |
| Glycerol | 1.098479 | -0.12798 | 0.170306 | 1.143889 | -1.22646 | -0.92817 | 0.04541 | 1.21E-05 | 7.28E-05 |
| Malic acid | 3.296901 | 3.407292 | 4.041762 | 3.492385 | 0.110391 | 0.744861 | 0.195484 | 1.83E-05 | 7.32E-05 |
| Myo-Inositol | 2.090478 | 1.45809 | 1.851852 | 1.432344 | -0.63239 | -0.23863 | -0.65813 | 1.66E-05 | 7.32E-05 |
| Aspartic acid | -5.2497 | -0.33153 | -4.02636 | -6.70497 | 4.918166 | 1.223338 | -1.45528 | 2.78E-05 | 9.52E-05 |
| Lactic acid | -2.1607 | -0.36896 | -0.33905 | -2.84353 | 1.791738 | 1.82165 | -0.68282 | 5.30E-05 | 0.000159 |
| Stearic acid | 0.050338 | -2.87551 | -2.46541 | -0.13792 | -2.92585 | -2.51574 | -0.18826 | 0.000104 | 0.000276 |
| Quinic acid | 3.06399 | 3.072054 | 3.326541 | 2.94019 | 0.008064 | 0.262551 | -0.1238 | 0.000225 | 0.000539 |
| Glucitol | 3.874245 | 4.149549 | 4.705752 | 4.189084 | 0.275303 | 0.831507 | 0.314838 | 0.000255 | 0.000555 |
| 1,3-Propanediol | 0.367169 | 0.475148 | 0.821102 | 0.157762 | 0.107979 | 0.453933 | -0.20941 | 0.000818 | 0.001636 |
| Ribofuranose | 1.103266 | 0.435249 | 1.561352 | 1.135569 | -0.66802 | 0.458086 | 0.032302 | 0.000974 | 0.001798 |
| Fructofuranose | 2.756699 | -0.69877 | 1.800167 | 2.762896 | -3.45547 | -0.95653 | 0.006197 | 0.002897 | 0.004966 |
| Arabinose | 1.707231 | 1.926177 | 1.773896 | 1.54241 | 0.218946 | 0.066665 | -0.16482 | 0.004328 | 0.006925 |
| Citric acid | -0.77701 | -1.16903 | -0.3551 | -0.55988 | -0.39202 | 0.421915 | 0.217139 | 0.007611 | 0.010745 |
| Fructose | 6.497367 | 6.372759 | 6.523849 | 6.259568 | -0.12461 | 0.026482 | -0.2378 | 0.007183 | 0.010745 |
| 1-Monopalmitin | 0.310616 | -0.62299 | -0.28023 | 0.055442 | -0.9336 | -0.59085 | -0.25517 | 0.008686 | 0.011582 |
| Urea | 1.44557 | 1.438246 | 1.811144 | 0.836559 | -0.00732 | 0.365573 | -0.60901 | 0.011697 | 0.014036 |
| Chlorogenic acid | -7.88915 | -6.14401 | -7.07468 | -6.68378 | 1.745143 | 0.81447 | 1.205371 | 0.011442 | 0.014036 |
| Fructopyranose | 0.789195 | 0.148297 | 1.116932 | -0.0004 | -0.6409 | 0.327737 | -0.7896 | 0.036222 | 0.041396 |

**Supplementary Table S3.** Receiver operating characteristic (ROC) statistics for all GC-MS identified metabolites detected in apple peel.

| **Metabolite** | **AUC** | **CI_lower** | **CI_upper** |
|---|---|---|---|
| Lupeol | 1 | 1 | 1 |
| Sucrose | 0.979167 | 0.875 | 1 |
| Ursolic acid | 0.958333 | 0.8125 | 1 |
| Mannose | 0.958333 | 0.8125 | 1 |
| 1-Monopalmitin | 0.9375 | 0.75 | 1 |
| Urea | 0.916667 | 0.75 | 1 |
| Chlorogenic acid | 0.854167 | 0.625 | 1 |
| Glucitol | 0.854167 | 0.625 | 1 |
| Lactitol | 0.833333 | 0.583333 | 1 |
| Glycerol | 0.833333 | 0.604167 | 1 |
| Gluconic acid | 0.8125 | 0.583333 | 1 |
| Ribofuranose | 0.791667 | 0.5 | 1 |
| Fucitol | 0.791667 | 0.458333 | 1 |
| Palmitic acid | 0.75 | 0.5 | 0.958333 |
| Fructofuranose | 0.708333 | 0.416667 | 1 |
| Oleanolic acid | 0.708333 | 0.4375 | 0.916667 |
| Myo-Inositol | 0.6875 | 0.416667 | 0.916667 |
| Glycerol monostearate | 0.666667 | 0.416667 | 0.916667 |
| Lactic acid | 0.666667 | 0.416667 | 0.916667 |
| Glucopyranose | 0.666667 | 0.395833 | 0.916667 |
| Allose | 0.666667 | 0.416667 | 0.916667 |
| Stearic acid | 0.645833 | 0.375 | 0.895833 |
| Citramalic acid | 0.645833 | 0.375 | 0.916667 |
| Quinic acid | 0.645833 | 0.333333 | 0.895833 |
| 1,3-Propanediol | 0.625 | 0.333333 | 0.875521 |
| Hydroxylamine | 0.625 | 0.333333 | 0.875521 |
| Malic acid | 0.625 | 0.291667 | 0.916667 |
| Mannitol | 0.604167 | 0.333333 | 0.854167 |
| Citric acid | 0.5625 | 0.208333 | 0.896354 |
| Fructopyranose | 0.5625 | 0.25 | 0.833333 |
| Fructose | 0.541667 | 0.25 | 0.833333 |
| Glucose | 0.541667 | 0.25 | 0.833333 |
| Arabinose | 0.479167 | 0.166667 | 0.833333 |
| Galactose | 0.458333 | 0.125 | 0.791667 |

**Supplementary Table S4.** Receiver operating characteristic (ROC) statistics for all GC-MS identified metabolites detected in apple pulp.

| **Metabolite** | **AUC** | **CI_lower** | **CI_upper** |
|---|---|---|---|
| Myo-Inositol | 1 | 1 | 1 |
| Chlorogenic acid | 0.9375 | 0.75 | 1 |
| Glucitol | 0.9375 | 0.791666667 | 1 |
| Threonine | 0.895833333 | 0.728645833 | 1 |
| 1-Monopalmitin | 0.895833333 | 0.708333333 | 1 |
| Glucose | 0.875 | 0.645833333 | 1 |
| Stearic acid | 0.854166667 | 0.625 | 1 |
| Fructofuranose | 0.854166667 | 0.625 | 1 |
| Malic acid | 0.833333333 | 0.541666667 | 1 |
| Glycerol | 0.833333333 | 0.604166667 | 1 |
| Erythronic acid | 0.770833333 | 0.5 | 0.979166667 |
| Mannitol | 0.770833333 | 0.375 | 1 |
| Fructose | 0.770833333 | 0.5 | 0.958333333 |
| Lactic acid | 0.729166667 | 0.458333333 | 0.9375 |
| 1,3-Propanediol | 0.6875 | 0.395833333 | 0.979166667 |
| Fucitol | 0.6875 | 0.416666667 | 0.916666667 |
| Sucrose | 0.666666667 | 0.416666667 | 0.916666667 |
| Citric acid | 0.645833333 | 0.375 | 0.875 |
| Fructopyranose | 0.645833333 | 0.375 | 0.895833333 |
| Urea | 0.625 | 0.333333333 | 0.875 |
| Aspartic acid | 0.583333333 | 0.333333333 | 0.833333333 |
| Arabinose | 0.5625 | 0.270833333 | 0.833333333 |
| Ribofuranose | 0.541666667 | 0.25 | 0.833333333 |
| Quinic acid | 0.520833333 | 0.25 | 0.791666667 |
